# Quasi epigenetic equilibrium: the implications of plasticity and variability on evolution and extinction

**DOI:** 10.1101/2025.08.27.672696

**Authors:** Puneeth Deraje, Matthew M. Osmond

**Affiliations:** Department of Ecology & Evolutionary Biology, University of Toronto

**Keywords:** Epigenetics, phenotypic plasticity, evolutionary rescue, phenotypic variability, mathematical model, population genetics

## Abstract

The phenotypic effects of epigenetic modifications, and thus their evolutionary consequences, depend on how the modifications interact with the underlying genetics and the surrounding environment. These interactions lead to a complex model that has so far prevented general analytical progress, and thus limited our understanding. Here, we show that the timescale difference between epigenetic and genetic changes create a quasi-epigenetic equilibrium (QEE). The QEE allows us to tackle the complexity of population epigenetic models by reducing them to their underlying population genetic models with effective parameters. Using this technique, we show how epigenetics modifies key evolutionary parameters, such as the strength of selection and dominance, which can have drastic evolutionary consequences on mutation-selection balance. Further, we show how the QEE allows us to analytically investigate the effect of epigenetics on the probability of a population escaping extinction in a harsh environment via adaptation – evolutionary rescue – by altering the number of potential rescue lineages and their probability of establishing. These calculations show that whether epigenetics helps or hurts population persistence depends non-trivially on the frequency and stability of epigenetic modifications.

## Introduction

An organism’s phenotype, and hence its fitness, depends not only on its genetic basis but on a multitude of non-genetic factors. These non-genetic factors can potentially alter the course of evolution, provided they are heritable (Day and Bonduriansky, 2011). An important source of such heritable non-genetic variation is epigenetics (Banta and Richards, 2018; Webster and Phillips, 2025). Epigenetics refers to a broad set of biochemical markers on DNA that can generate distinct phenotypes from the same genetic code by a variety of mechanisms, such as altering gene expression, alternative splicing, and modulating chromatin accessibility (Aristizabal *et al*., 2020). While these mechanisms are themselves important for the growth and development of an organism, they can also be a source of heritable variation infitness that influences the course of evolution (Kronholm, 2023).

The role of epigenetics in evolution is the focus of a growing body of theoretical research (Geoghegan and Spencer, 2012, 2013a,b; Kronholm and Collins, 2016; Slatkin, 2009; Stenøien and Pedersen, 2005; Webster and Phillips, 2024). Most of these models focus on either phenotypic plasticity (Chernomas and Griswold, 2024; Gunnarsson *et al*., 2020; Lambert *et al*., 2025) – the ability of a genotype to produce different phenotypes in different environments - or phenotypic variability (Carja and Plotkin, 2017, 2019) – the ability of a genotype to exhibit different phenotypes in the same environment. However, plasticity and variability do not exist in isolation, i.e., epigenetic modification can alter phenotypes to different degrees in different environments (Feiner *et al*., 2021). To better understand the role of epigenetics on evolution we need models that allow this interaction between epigenetics and environment.

Incorporating epigenetics into evolutionary models increases the number of variables in the model (one for each epigenetic variant). As a result, many models allow only one allele to be modified, resort to simulations, and/or restrict their analysis to specific parameter regimes. For example, Kronholm and Collins (2016) built a model that allows multiple alleles to be modified but restricted the analysis to simulations where epigenetic modification rates and phenotypic effects are independent of the underlying genotype. And, more recently, Webster and Phillips (2024) considered a model that allows for epigenetic variants of two alleles but were analytically limited to special cases, such as neutrality or additivity. This lack of tractability poses a key challenge to understanding the effect of interactions between epigenetics and genetics and extending to more complex epigenetic architectures. For example, multiple genes in a gene-regulatory network could each undergo epigenetic changes (Roy and Kundu, 2014) or there could be more complex mechanisms of epigenetic modification, such as paramutation (Hollick, 2017), which only occurs in heterozygotes. A general analysis of realistically-complex epigenetic models requires mathematical techniques that can tackle their high dimensionality.

A key source of interest in epigenetics is its potential to help populations survive harsh changes in the environment (O’Dea *et al*., 2016). For example, epigenetic mechanisms have been implicated in a wide variety of scenarios, including resistance to common antibiotics (Ghosh *et al*., 2020), drug resistance in cancer (Karami Fath *et al*., 2022), arthropod pest resistance (Mogilicherla and Roy, 2023), insecticide resistance (Oppold and Müller, 2017), plant-pathogen interactions (Boyko and Kovalchuk, 2011), tolerance to heat (Jablonka and Raz, 2009), and adaptation to temperature (Yaish *et al*., 2011). However, epigenetic changes need not always be beneficial. For instance, when exposed to certain harsh environments,*Daphnia*exhibit epigenetic responses that lower theirfitness (Shahmohamadloo *et al*., 2025). In addition, epigenetics can potentially hinder genetic evolution by reducing the relativefitness of otherwise beneficial genetic mutants (O’Dea *et al*., 2016). It is therefore not clear how epigenetics affects the probability of evolutionary rescue (O’Dea *et al*., 2016), i.e., the probability that a population escapes extinction in a novel environment by evolution.

Here, we examine a very general one-locus population epigenetic model. We assume there are two genetic alleles, each with an epigenetic variant with arbitrary fitness, and consider both haploid and diploid selection. The rate at which each allele is modified and the rate at which its modifications are reset are also arbitrary. By exploiting the relatively fast epigenetic dynamics, we demonstrate the existence of a quasi equilibrium between a genetic allele and its epigenetic variant, allowing us to reduce the analytical complexity of the problem. We use this tractability to explore the effect of epigenetics on mutation-selection balance, extend to a more complex form of epimutation (paramutation) and address a key biological question: how does epigenetics affect evolutionary rescue?

## Model

Consider a panmictic population of organisms or cells. There are two genetic alleles, the wildtype *A* and the mutant *a*. Each generation, allele *A* can be epigenetically modified to *B* with probability *m*_*A*_ and allele *a*can be epigenetically modified to *b* with probability *m*_*a*_. We refer to this as standard epimutation and later consider a more complicated form of epimutation known as paramutation. These modifications are reversed (reset) each generation with probability *t*_*A*_ and *t*_*a*_ respectively. For clarity, we will refer to alleles that are not epigenetically modified (*A* and *a*) as unmodified alleles, while genetic alleles will refer to the genetic allele irrespective of the epigenetic marker (e.g., genetic allele *A* includes both unmodified *A* and modified *B* forms). Genetic mutation between unmodified *A* and *a* occurs with probability *µ* every generation.

Further, since epigenetic marks can alter genetic mutation rates (Gonzalgo and Jones, 1997; Habig *et al*., 2021), we allow genetic mutation between the epigenetic variants *b* and *B* to occur with a different probability,*ν*. See Fig. 1 (left) for a visual depiction of the transition rates between epialleles.

**Figure 1.**
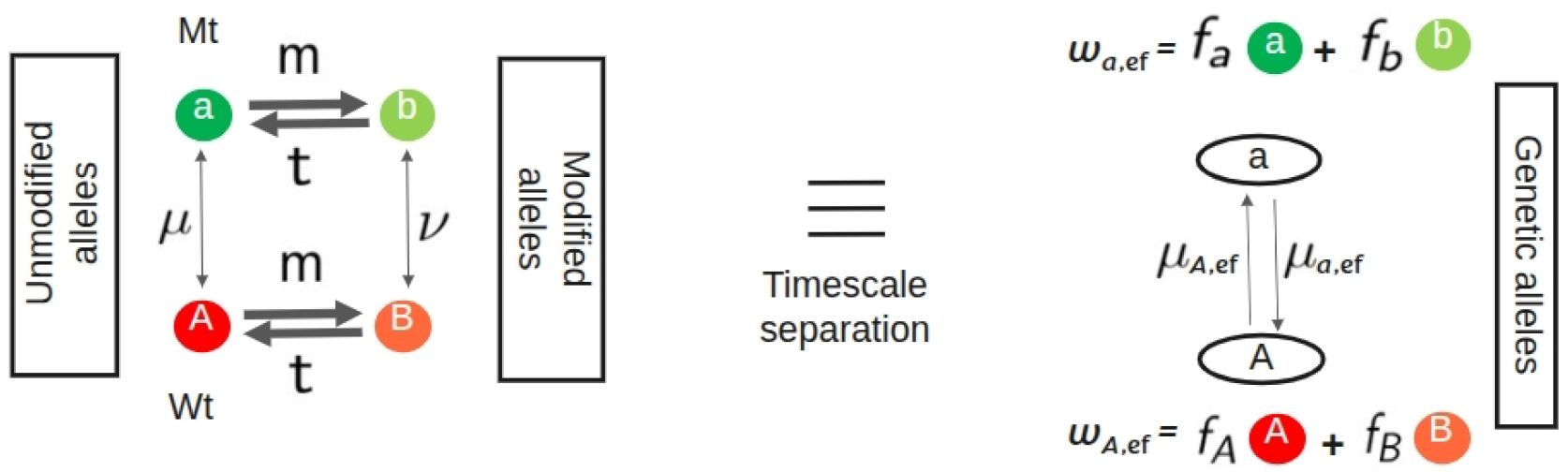
Quasi epigenetic equilibirum (QEE). (left) The full four epiallele system. (right) The underlying two genetic allele system. The rapidity of the epigenetic dynamics relative to the mutation-selection dynamics creates a QEE (*f*_*x*_) that reduces the four-allele system to a two-allele system with effective mutation rates and fitnesses.

To be clear, in our model *m*_*x*_ is the probability of an offspring acquiring an epigenetic modification their parent did not have and, similarly, *t*_*x*_ is the probability of an offspring losing a modification their parent did have. It is tempting to think of *t*_*x*_ as the probability of losing a modification during epigenetic reprogramming in gametogenesis (Feng *et al*., 2010), which we can define as *t*_*x*,bio_. However, the offspring can reacquire the modification after gametogenesis. Therefore, the reset probability can be understood as *t*_*x*_ = *t*_*x*,bio_(1−*m*_*x*_), the probability of losing the modification during gametogenesis and not reacquiring it.

We first consider the case where all (epi)genotypes have absolute fitness above replacement so that the evolutionary dynamics are completely characterized by relativefitness. We set the wildtype allele/homozygote relative fitness to 1 and write the mutant allele/homozygote relative fitness as 1−*s*. We make no assumptions about the relative fitness of any (epi)genotype in developing the theory. Our key assumption is that mutation is weaker than selection and that selection is weaker than epigenetic modification and reset (*µ*,*ν~*𝒪(*ϵ* ^2^);*s~O*(*ϵ*);*m* _*x*_, *t*_*x*_ *~*𝒪(1)). Note that this allows for guaranteed reset, *t* _*x*_ = 1, which corresponds to no epigenetic inheritance.

In our model, we associate each individual with a single epigenotype but, of course, epigenetic markers can differ between cells in multicellular individuals. Our model accounts for this when individuals initiated with a given epigenotype have the same fitness, e.g., when they give rise to a similar pattern of epigenotypes across cells.

### Quasi epigenetic equilibrium

We begin by formalizing a technique to analyze epigenetic systems in an analytically tractable and biologically informative way. This uses the key observation that epigenetic rates are generally much faster than the mutation-selection dynamics (Rando and Verstrepen, 2007). Previous theoretical work has shown that the distribution of epigenetic states at equilibrium is relatively insensitive to mutation-selection dynamics in empirically relevant parameter regions (Slatkin, 2009; Webster and Phillips, 2024). Here we show that this is true not just at equilibrium. The distribution of epigenetic states of a given genetic allele can be perturbed by mutation and selection but reaches a new equilibrium almost instantaneously. We call this the quasi epigenetic equilibrium (QEE) in analogy to the quasi linkage equilibrium (QLE), where the effects of recombination occur faster than mutation and selection (Kimura, 1965). Here the QEE allows us to reduce our four epiallele system to classical two allele systems with effective fitnesses and mutation probabilities that depend on the QEE (Fig. 1, right). This allows us to use extensive existing population genetic results to gain insights into the effects of epigenetics on evolution.

We first exemplify how to obtain the QEE and demonstrate its accuracy under haploid selection. We then examine diploid selection and a more complex form of epimutation (paramutation). Finally, we explore the implications for evolutionary rescue.

### Haploid selection

Under haploid selection, the unmodified wildtype allele, *A*, has relative fitness *w*_*A*_ = 1 and the unmodified mutant allele, *a*, is deleterious with relative fitness *w*_*a*_ = 1 − *s*. The fitnesses of the epigenetically modified alleles, *x* = *a, b*, are denoted *w*_*x*_ = 1 −*h*_*x*_*s*. If *h*_*x*_ = 0, the respective allele has fitness identical to the unmodified wildtype *A*, while if *h*_*x*_ = 1, it has fitness identical to the unmodified mutant *a*. We make no assumptions about the range of *h*_*x*_.

The dynamics of the epiallele frequencies, *p*_*x*_, are given by

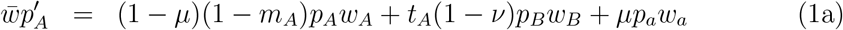

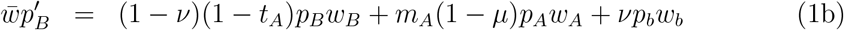

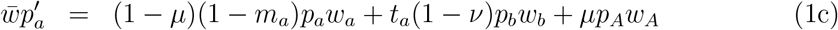

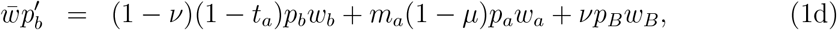

Where 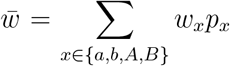 is population mean fitness. Given the four variables sum to one we can rewrite these recursions in terms of the frequency of the genetic mutant allele *a* irrespective of its epigenetic states, *p*_*a*,gen_ = *p*_*a*_ +*p*_*b*_ = 1−*p*_*A*,gen_, and the relative frequency of the epigenetic states for each genetic allele, *f*_*b*_ = *p*_*b*_/*p*_*a*,gen_ = 1 −*f*_*a*_ and *f*_*B*_ = *p*_*B*_*/p*_*A*,gen_ = 1 −*f*_*A*_. The recursions for these are

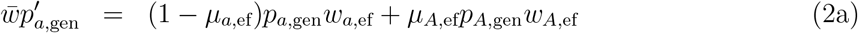

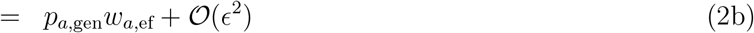

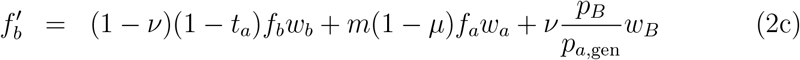

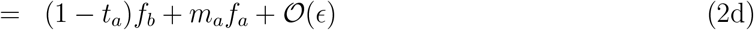

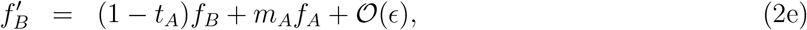

Where

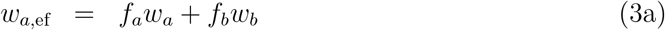

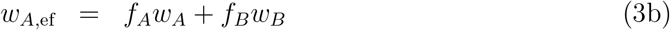

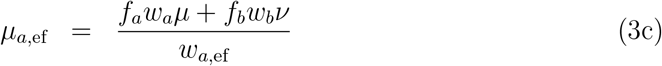

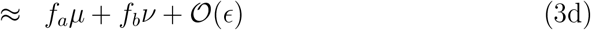

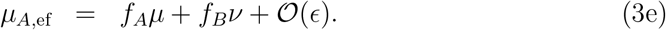

We refer to *w*_*x*,ef_ and *µ*_*x*,ef_ as the effective fitness and effective mutation rate of genetic allele *x* (i.e., averaged over epigenetic states).

Note that the leading term in the change in allele frequency, 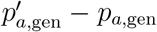, is of 𝒪 (*ϵ*) but the leading term in the change in the proportion modified, 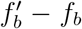, is of 𝒪 (1). Therefore, *f*_*b*_ equilibrates faster than *p*_*a*,gen_ and hence genetic allele frequency can be considered constant on the time scale of the epimutation dynamics. We call the equilibrium for *f*_*b*_, given a constant *p*_*a*,gen_, the quasi epigenetic equilibrium (QEE). The QEE can be perturbed by changes in genetic allele frequency but quickly reaches a new equilibrium. In this particular case, the QEE up to leading (zeroth) order (in *ϵ*) is independent of the genetic allele frequency,

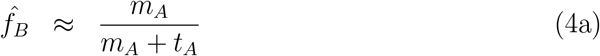

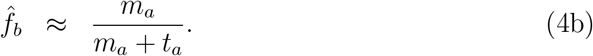

As these quantities will appear frequently, we define 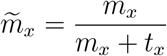 and 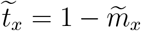. Fig 2a shows that this leading order QEE is accurate and reached quickly, typically within *~* 10 generations. The first-order correction is given in Section S1.

**Figure 2.**
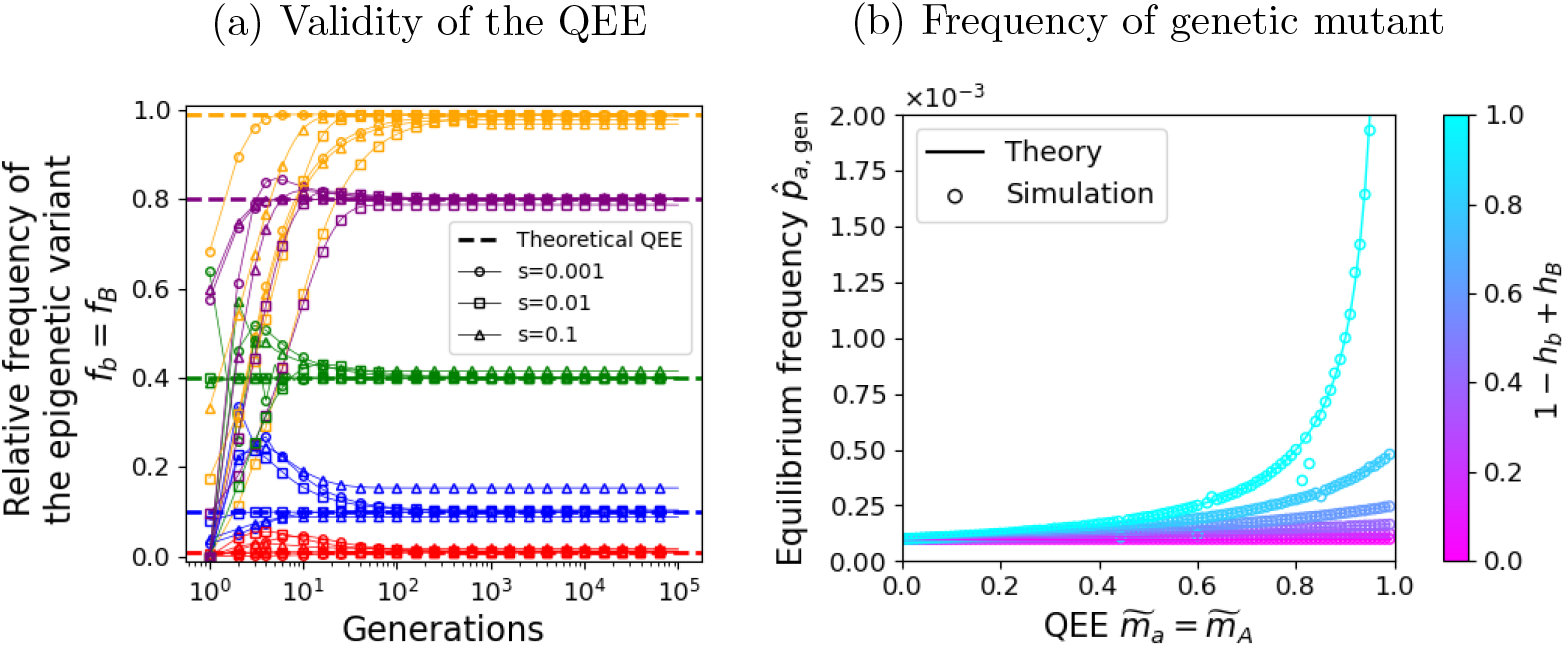
Validity of the QEE its consequences under haploid selection. (a) The proportion of genetic alleles *A* and *a* that are epigenetically modified (*f*_*B*_ and *f*_*b*_, respectively) for different strengths of selection (different shapes) and different epimutation/reset rates (different colors) through time. The first order QEE approximation (dashed lines, Eq. 4a and 4b) is reached within a few generations even for relatively strong selection. (b) The frequency of the genetic mutant *a* as a function of the QEE for different fitness of the alleles (different colors). Circles are obtained through deterministic simulations after 10^5^ generations starting with only the wildtype allele *A* and lines are our analytical approximation (Eq. 5). For each QEE value, the specific epimutation and reset probabilities were chosen randomly such that 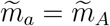. Other parameters: *µ* = *ν* = 10^−6^, *s* = 0.01

There are two key consequences of this form of the QEE. First, the effect of epigenetics on genetic evolution is completely determined by the ratio of epigenetic rates (modification and reset). The absolute values of the rates are irrelevant. Therefore, the same evolutionary effect could occur in two ways: *f*_*x*_ can be large either with high modification probability irrespective of reset or low modification with stable transgenerational inheritance. An example of the former is a gene near a transposable element (TE), which can be repressed consistently due to efficient silencing in organisms with extensive reset machinery (Choi and Lee, 2020). The latter is exemplified by gene regulation via transgenerationally inherited sRNA in *C*.*elegans* (Rechavi and Lev, 2017).

The second consequence of this form of QEE arises because it is frequency-independent. When we plug the leading order QEE into the effective fitnesses, *w*_*x*,ef_, and mutation rates, *µ*_*x*,ef_, they are impervious to allele frequency and the dynamics of the genetic alleles (irrespective of their epigenetic markers) reduce to those of the classical 2-allele model. The myriad of results for this well-known model can then be immediately applied. For example, the equilibrium allele frequency is

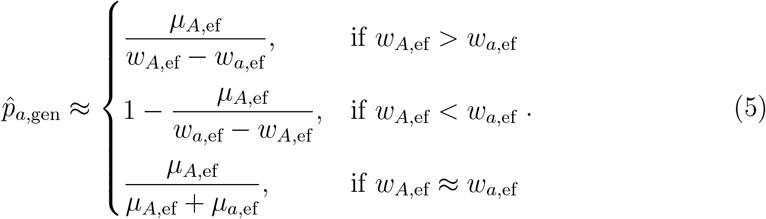

An interesting consequence of this is that epigenetic modifications can increase the frequency of mutants at equilibrium both by reducing the effective fitness of the wildtype allele (which occurs when modification reduces wildtype fitness, *h*_*B*_ *>* 0) and by increasing the effective fitness of the mutant (which occurs when modification increases mutant fitness, *h*_*b*_ *<* 1) (Fig 2b).

### Diploid selection

Now considering diploid selection, the relative fitness of the unmodified wildtype homozygote is *w*_*AA*_ = 1, the relative fitness of the unmodified mutant homozygote is *w*_*aa*_ = 1 −*s*, and the relative fitnesses of all other genotypes *xy* are written *w*_*xy*_ = 1 −*h*_*xy*_*s*.

The leading order QEE, 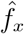, and the resulting effective mutation rates, *µ*_*x*,ef_, are identical to the haploid case (equations 4a, 4b, and 3c-e). The only difference is that now we need to calculate the effective fitnesses of diploid genotypes,

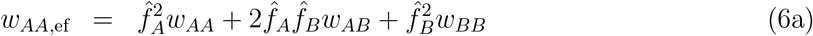

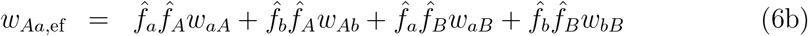

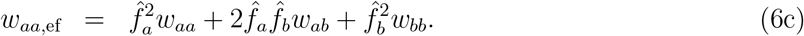

With this conversion, the dynamics of the genetic alleles under the QEE reduce to the classical 2-allele diploid model. In particular, the equilibrium frequency of the genetic mutant is

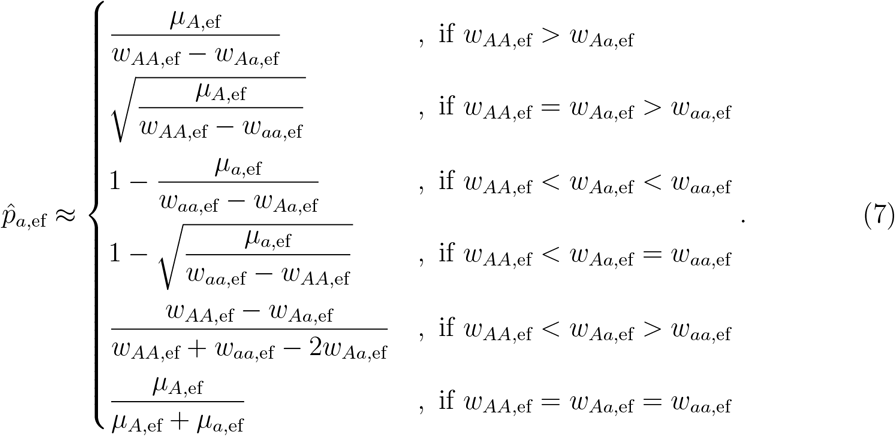

An interesting consequence of the effective fitnesses depending on the QEE is that the dominance coefficient of the genetic mutant is sensitive to the epimutation rates. As a result, small changes in the epimutation rates can lead to drastic changes in genetic allele frequencies at equilibrium (Fig. 3a). Such effects are most prominent if diploid epigenotypes with a single modified allele have higher (or lower) fitness than completely unmodified or modified epigenotypes (Fig. 3b) but can arise in a wide variety of situations.

**Figure 3.**
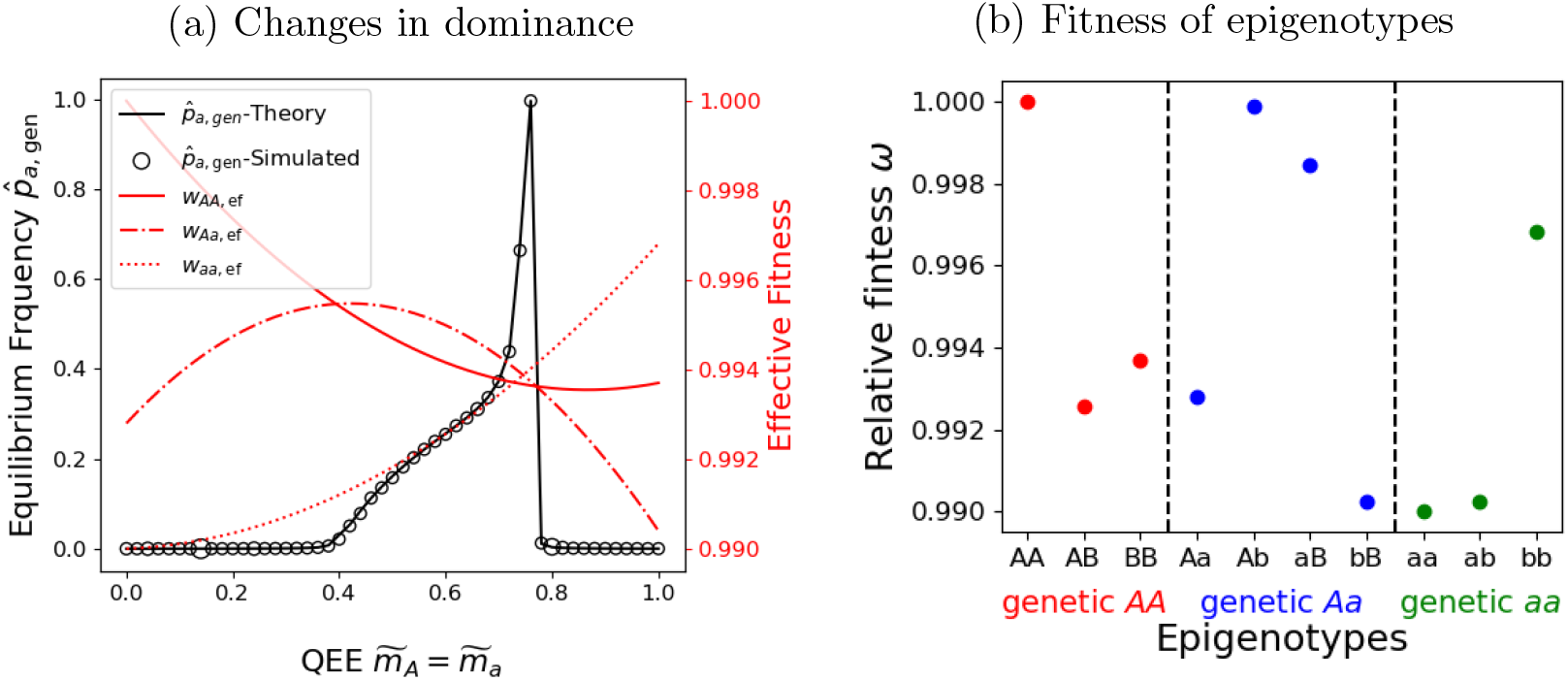
The QEE influences dominance and the consequent equilibrium under diploid selection. (a) The equilibrium frequency (black axis) of the genetic mutant allele *a* from full simulations (empty circles) and analytical approximations (solid line, Eq. 7) as a function of the QEE, 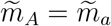. Also plotted are the effective fitnesses (red axis). The values of *m*_*x*_ and *t*_*x*_ are chosen randomly for each simulated 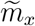. (b) The relative fitness of each epigenotype is colored by the effective genotype it contributes to. Within each color, the epigenotypes are arranged such that higher values of 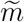 lead to a higher proportion of epigenotypes on the right. Other parameter values: *µ* = *ν* = 10^−6^.

### Paramutation

To demonstrate the utility of the QEE in more complex scenarios, we now consider paramutation. Paramutation, observed in several organisms such as maize, tomato, *Arabidopsis*, a plant pathogen, mice, fungus and *Drosophila* (Chapter 1 of Haring *et al*., 2005), is an epimutation that occurs only in heterozygotes. Only one of the unmodified alleles, *A* or *a*, can be epigenetically modified. We call this allele paramutable. The other allele can induce epigenetic modifications on the paramutable allele when present together in a heterozygote. We call this allele paramutagenic. Further, the epigenetically modified allele can itself modify the paramutable allele when in a (epi)heterozygote (Geoghegan and Spencer, 2013a). Here we ignore standard epimutation to focus on the unique effects of paramutation.

The fact that paramutation only occurs in heterozygotes makes it frequencydependent. Despite this, the separation of timescales still works but now the leading order QEE depends on the genetic allele frequencies. In particular,

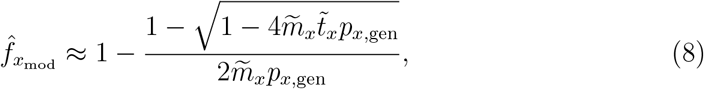

where *x* is the paramutable allele (*a* or *A*) and *x*_mod_ is its modified form (*b* or *B*). Plugging this into the effective diploid fitnesses (Eq. 6), the dependence of the QEE on allele frequency means the genetic alleles are under frequency-dependent selection. A general analysis of frequency-dependent selection, even in the absence of epigenetic dynamics, is usually challenging (Cockerham *et al*., 1972; Otto *et al*., 2008). However, the QEE simplifies when the paramutable allele is sufficiently rare or common. We divide the analysis into these two cases.

When the mutant, *a*, is paramutable (so the (epi)alleles are *A, a*, and *b*), it is present at low frequency and nearly always found in a heterozygote, *Aa*. Therefore every mutant allele undergoes epimutation with probability *m*_*a*_ and the QEE reduces back to 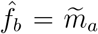. To see this, take the Taylor series of 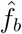 (Eq. 8) around *p*_*a*,ef_ = 0, which will give 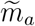 to leading order. This allows us to ignore the frequency dependence and we can use equilibrium allele frequencies from the diploid case with standard epimutation (Eq. 7, see Section S2 for a special case). The approximation works very well (Fig. 4a-c, Fig. S2).

**Figure 4.**
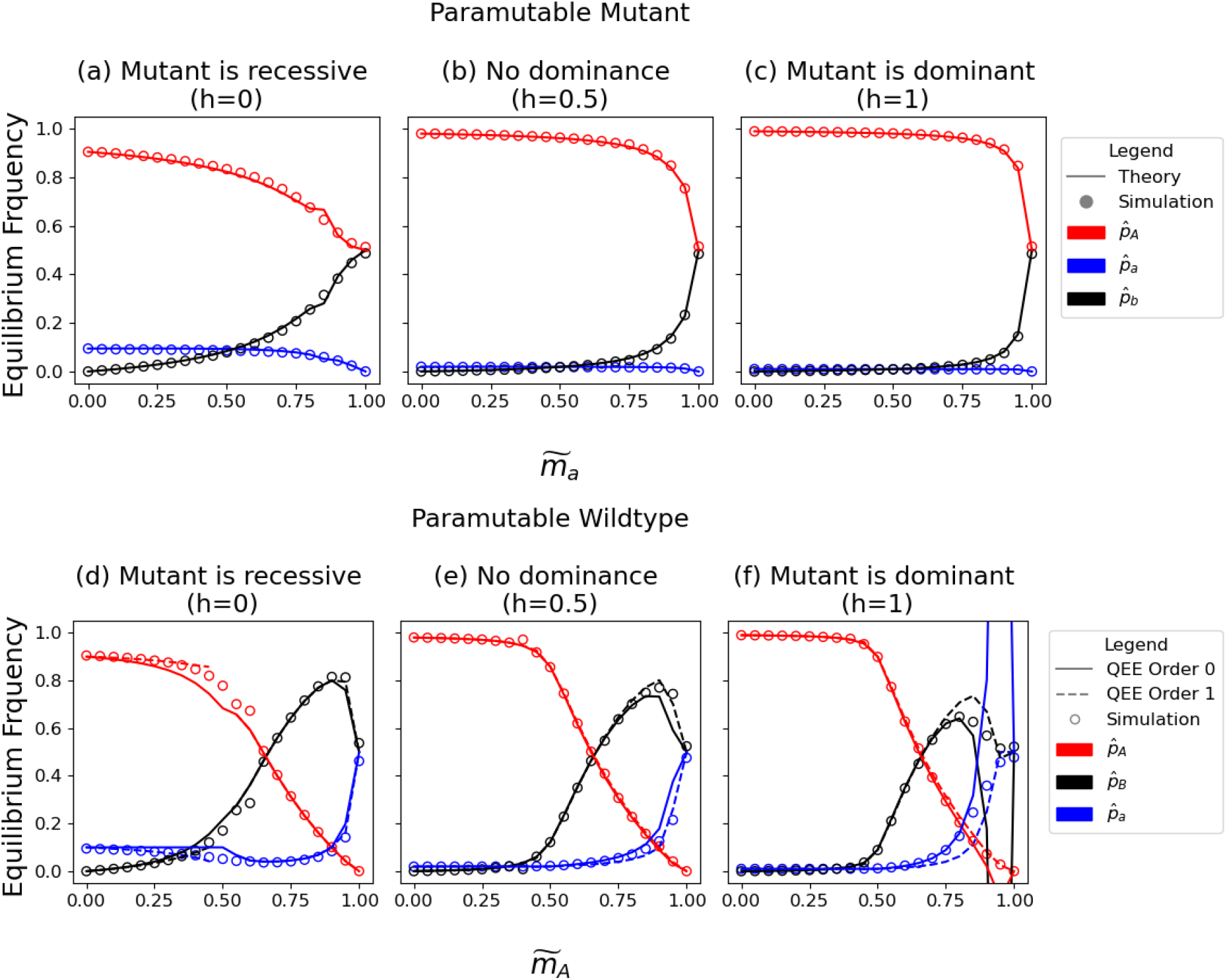
The effect of paramutation on equilibrium allele frequencies. (a-c) The mutant allele *a* is paramutable and the epiallele *b* has the same fitness as the wildtype *A*: *h*_*AA*_ = *h*_*Ab*_ = *h*_*bb*_ = 0 and *h*_*Aa*_ = *h*_*ab*_ = *h*. (d-f) The wildtype allele *A* is paramutable and the epiallele *B* has the same fitness as the mutant *a*: *h*_*aa*_ = *h*_*aB*_ = *h*_*BB*_ = 1 and *h*_*Aa*_ = *h*_*AB*_ = *h*. In all figures the empty circles are from deterministic simulations while the lines are analytical approximations. Solid lines use the zeroth order QEE: (a) Eq. S2, (b-c) Eq. 7, (d-f) Eq. 7 but with 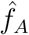 from Eq. 9. The dashed lines in (d-f) use the QEE up to first order: Eq. S4 and Eq. S5. Other parameters: *µ* = *ν* = 10^−4^, *s* = 0.01.

It is more complicated when the unmodified paramutable allele is the wildtype *A* (meaning we have epialleles *A, a*, and *B*). This is because, in addition to heterozygotes (*Aa*), the paramutable allele is now often found in homozygotes (*AA*), where it cannot be modified. The frequency dependence in the QEE then becomes important. Specifically, a sufficient frequency of paramutagens, *a* and *B*, is required for paramutation to have an effect and once it does the frequency of paramutagens can grow quickly (i.e., there is a positive feedback). Despite the importance of frequencydependence in this case, we can still simplify the problem by separating the QEE into two regimes. Taking a Taylor series of 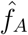(Eq. 8) around *p*_*a*,ef_ = 0 to leading order, we have

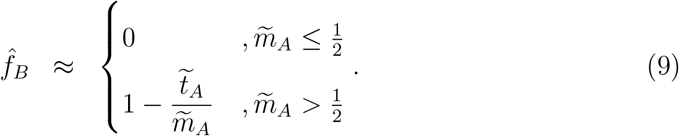

From this we see that frequency dependence makes the QEE more complicated (compare to Eq. 4a). In particular, the QEE is now a stepwise function because of the positive feedback, and the frequency of paramutagens only escalates when paramu-tation is effectively larger than reset, 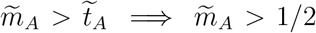. Despite the more complex form, the leading-order approximation is still frequency-independent. This frequency-independent leading order approximation can be used to obtain the equilibrium frequency of the genetic alleles (Eq. 7), which does a reasonable job (solid lines in Fig. 4d-f and Fig. S1).

With a paramutable wildtype there are rare cases where the frequency dependence cannot be ignored. In these cases we need to use the first-order term of the QEE (Eq. S3), which depends on the frequency of genetic alleles. It is then not possible to obtain analytical approximations in general, but they can be attained case by case, as demonstrated in Section S2.

### Evolutionary rescue

Epigenetic modification is hypothesized to play an important role in population persistence during harsh environmental changes (O’Dea *et al*., 2016). Here, we show how the QEE can be used to formally explore the role of epigenetics in population survival after an abrupt environmental change. As in the above analyses, the QEE allows us to gain insight from existing population genetic theory.

At the time of environmental change we assume the population is at the equilibrium computed in the previous sections and has a population size *N*_0_. After the environmental change, the wildtype allele/homozygote has absolute fitness 1 −*r* < 1, potentially putting the population at risk of extinction. However, the mutant allele/homozygote has absolute fitness 1 −*r* + *s*′ *>* 1 and so could potentially rescue the population. The remaining (epi)alleles/(epi)genotypes have absolute fitness 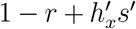. As typical for evolutionay rescue, we treat the post-change dynamics as a branching process. We allow the modification rates to change with the environment and hold mutation rates constant. Figure 5 gives a visual overview of the model.

**Figure 5.**
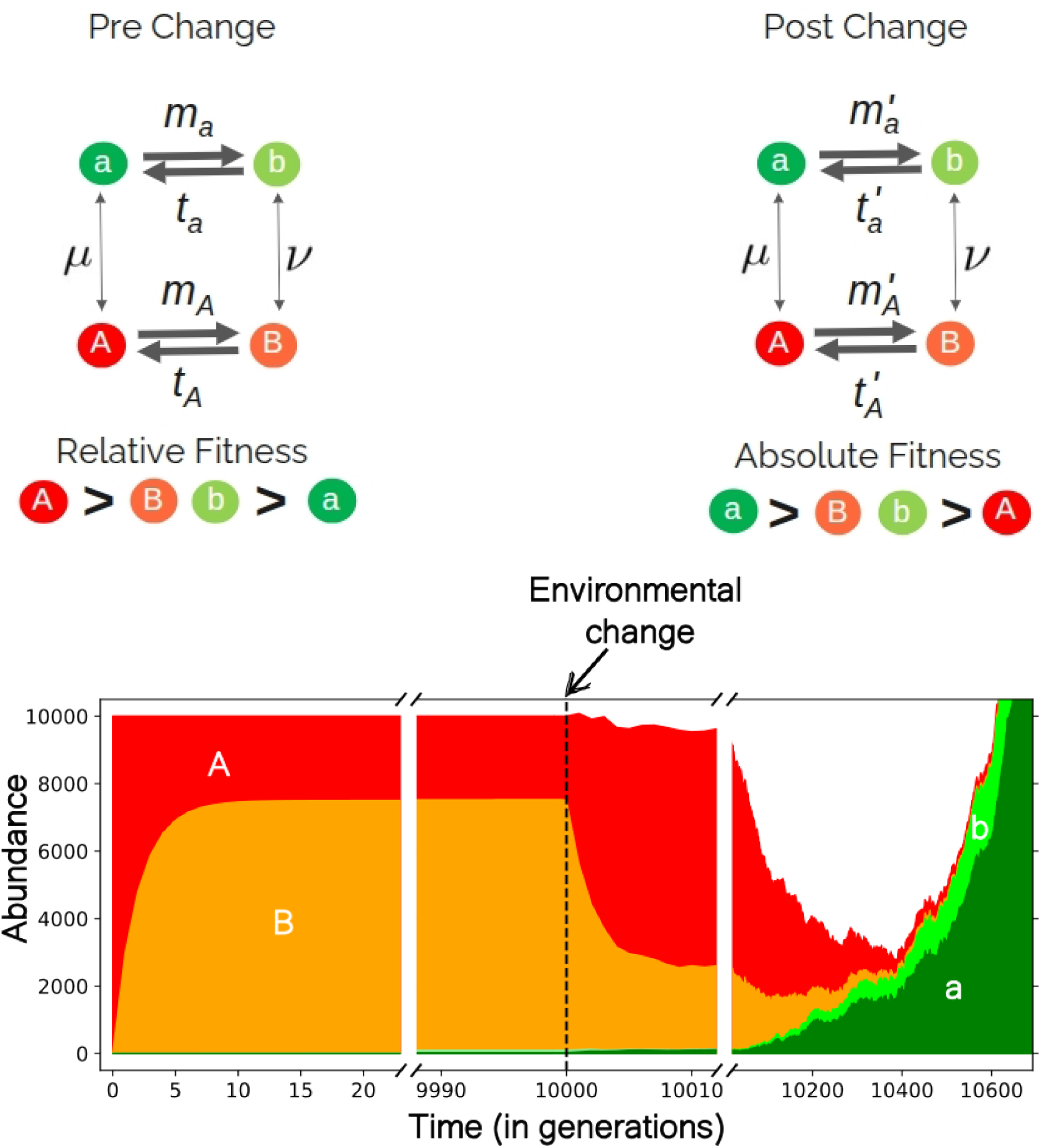
A model of evolutionary rescue with epigenetics. (top) The epigenetic modification rates, mutation rates, and fitnesses before (left) and after (right) the environmental change. (bottom) An example of the population dynamics before and after the environmental change (dashed line). Before the environmental change, the epiallele frequency dynamics are determined by relative fitness as in previous sections. After the environmental change, we treat the number of each type of epialleles as a branching process. Note the rapid (within 10 generations) equilibration of epigenetic dynamics compared to genetic dynamics, both at the beginning and after the environmental change.

Evolutionary rescue at a single locus without epigenetics has been studied for haploids (Orr and Unckless, 2008, 2014) and for diploids (Glémin and Ronfort, 2013; Uecker, 2017). We first outline the basic theory of those models, which we use and compare to below. The probability of rescue without epigenetics depends on three key components: (a) the number of mutants in the standing genetic variation, *N*_sgv_, (b) the number of mutants generated *de novo* during the wildtypes’ decline, *N*_dnm_, and (c) the establishment probability of a single mutant, *p*_est_. The probability of rescue is then given by

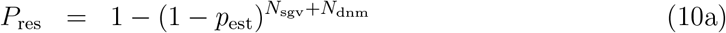

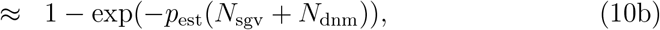

which is the probability that at least one of the mutants establishes.

Epigenetic variation can affect each of the three key components to increase or decrease the probability of rescue by the establishment of genetic mutants in complex ways. It also provides a new avenue for rescue mediated by the epigenetic variant of the wildtype, which we next outline.

### Haploid selection

#### Evolutionary rescue by epigenetics alone

To understand the baseline effect of epigenetic variation, we first assume there is no genetic variation, i.e., there is no genetic mutant. The survival of the population is then completely contingent on the epigenetic variant of the wildtype. The effective absolute fitness of the genetic wildtype allele post-change is 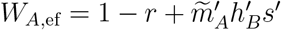, where primes denote values in the new environment. Its establishment probability is then roughly *p*_est,*A*_ = 2(*W*_*A*,ef_ − 1) when *W*_*A*,ef_ *>* 1 and otherwise 0 (Haldane, 1927). Establishment is only possible when *W*_*A*,ef_ *>* 1, which requires a sufficiently large epimutation probability, 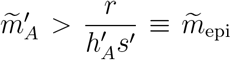. We call this the epigenetic threshold, below which rescue by epigenetics alone is not possible. The probability of rescue by epigenetics alone can then be approximated by

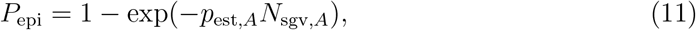

where

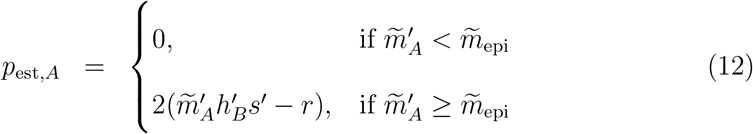

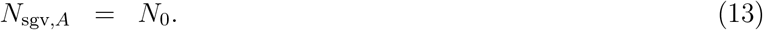

The standing epigenetic variation is irrelevant because the new QEE is reached rapidly (Fig. 5) and therefore completely determines the population’s fate.

Given a large initial population size, *N*_0_ *>>* 1, survival is either very unlikely or almost guaranteed, except for a very tiny region close to 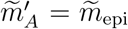 (Fig. 6a). This is because the rapid transitions between the wildtype and the epiallele make them behave like a single entity, effectively making every lineage subcritical (*W*_*A*,ef_ *<* 1) or supercritical (*W*_*A*,ef_ *>* 1). Near criticality, in addition to the zeroth-order QEE, the absolute value of modification rates, 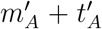, also has an effect. Since the rescue probability is zero at criticality, minor deviations in the QEE due to the first-order terms can have a major effect. For instance, at 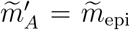, the difference in rescue probability between 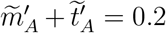 and 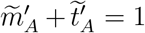 in Fig. 6a are due to a difference of 0.0002 in the effective fitness of *A* caused by a 0.02 change in the QEE (Fig. 6b and Section S3).

**Figure 6.**
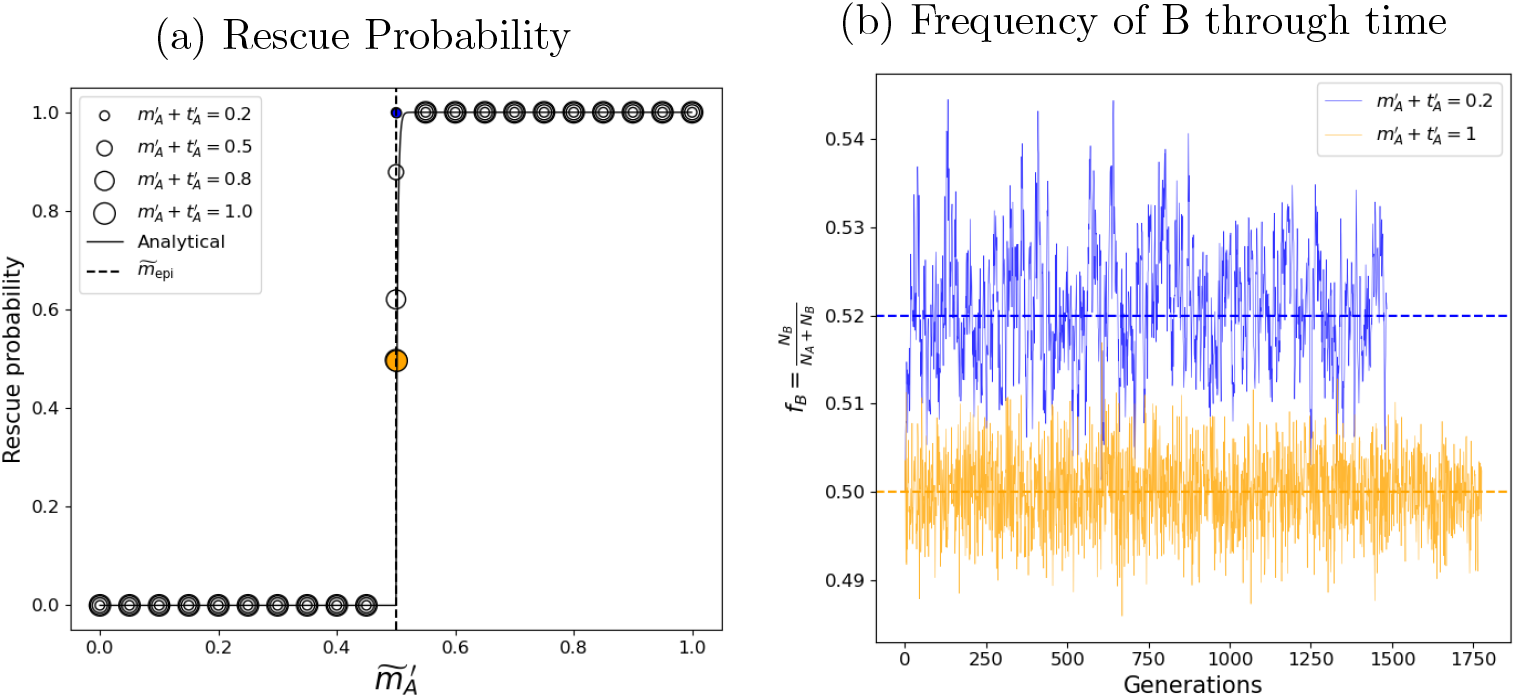
Evolutionary rescue by epigenetics alone. A haploid population’s response to environmental change when survival is contingent solely on an epigenetic variant, i.e., there are no genetic mutants. (a) Rescue probability as a function of the QEE in the new environment, 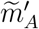. The solid line is the theoretical prediction (Eq. 11), the vertical line indicates criticality (*W*_*A*,ef_ = 1), empty circles are simulation results (1000 replicates). (b) The frequency of*B*for different values of 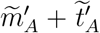 (different colors) when the zeroth order QEE leads to criticality 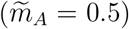. The dashed lines are the first-order QEE (Eq. S6) Here*m* _*A*_ =*t* _*A*_ = 0.8 and 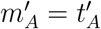. Other parameters: *r*= 0.01, *s*= 0.01, *s* ′ = 0.02, 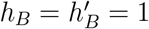.

#### Interactions between epigenetics, genetics, and the environment

To understand the consequences of interactions between epigenetics, genetics, and the environment for evolutionary rescue we now return to the full model with a genetic mutant and its epigenetic variant. Rescue can now occur either due to the epigenetic variant of the wildtype (as above), the genetic mutant, or both.

As seen in the pure epigenetic case, the epigenetic variant of the wildtype can rescue the population (irrespective of the genetic mutant) when the effective rate of epimutation is greater than the epigenetic threshold. The probability of rescue contingent purely on the wildtype epigenetic variant, *P*_epi_, is given by Eq. 11 with 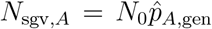, the number of genetic wildtype alleles in the standing variation (computed using Eq. 5).

The effective absolute fitness of the genetic mutant is 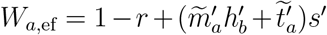. Its effective establishment probability is then roughly *p*_est,*a*_ = 2(*W*_*a*,ef_−1) when *W*_*a*,ef_ *>*1 and otherwise 0 and so cannot establish when 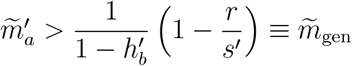, which we call the genetic threshold. The probability of rescue due to the establishment of the genetic mutant is then given by

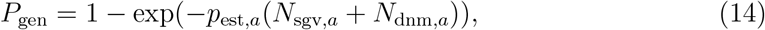

where

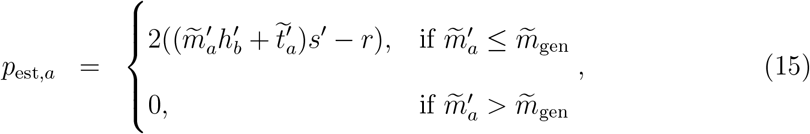

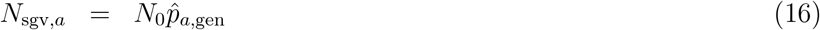

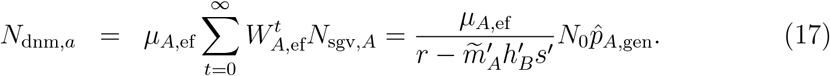

From this we see that epigenetic modification, characterized by 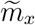 and 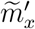, affects the probability of rescue by the genetic mutant in multiple ways. By altering effective fitnesses and mutation rates, modification prior to environmental change affects the standing variation (*N*_sgv,*a*_) while modification in the new environment affects the number of mutants that arise *de novo* (*N*_dnm_) and the establishment probability of the mutant, *p*_est,*a*_.

Putting the two sources of survival together, the total probability of rescue is

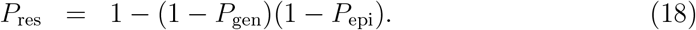

Whether epigenetics increases or decreases the rescue probability depends on the fitness of the epigenetic variants and the correlation of the epigenetic rates between alleles and environments.

To more concretely demonstrate the possible implications of epigenetics on survival, we now assume the fitness of each epigenetic variant is intermediate between the genetic wildtype and mutant 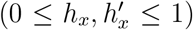 and focus on three scenarios: (1) epimutations only increase fitness, (2) modification and reset probabilities are allelespecific but not environment-specific, and (3) modification and reset probabilities are environment-specific but not allele-specific.

Given that epigenetic variants have intermediate fitness, epimutations happen only in the beneficial direction when only the mutants undergo epimutation pre-change, 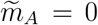, and only the wildtypes undergo epimutation post-change, 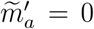. This is the case that is usually posited when claiming epigenetics aids survival in a new environment (Lambert *et al*., 2025; O’Dea *et al*., 2016). As expected, the probability of rescue is always higher with than without epimutations (Figure 7a). Further, increasing either of the non-zero effective epimutation rates increases the survival probability. Specifically, increasing the epimutation rate of the wildtype post-change reduces the decline rate of the wildtype, increasing the number of *de-novo* mutants (Fig. S3a), and at high enough rates 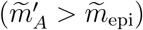 can lead to survival via epigenetics alone. Increasing the epimutation rate of the mutant pre-change increases the number of mutants in the standing (epi)genetic variation (Fig. S4a).

**Figure 7.**
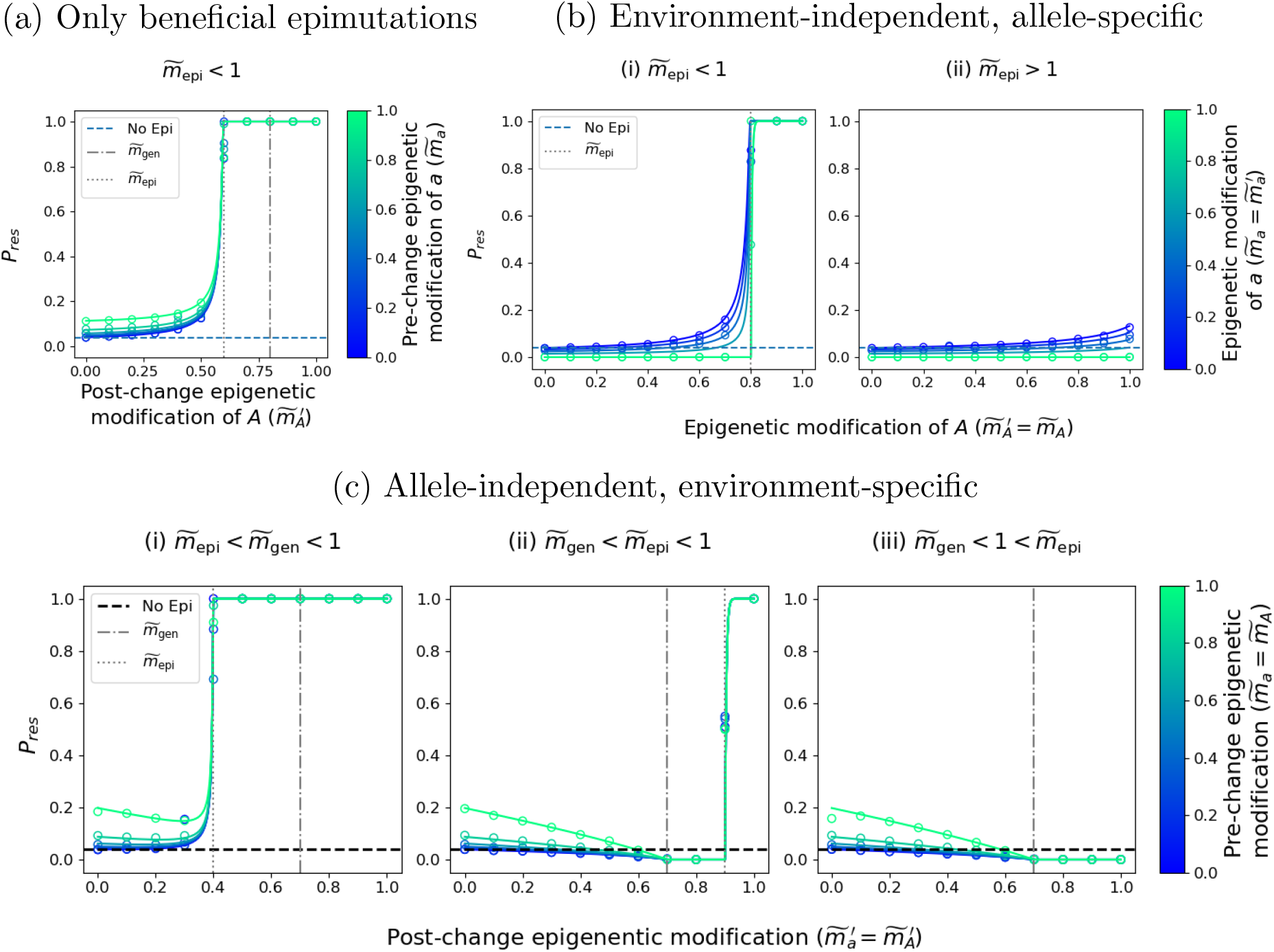
The probability of evolutionary rescue with epigenetics in haploids. Rescue probabilities for the haploid model (Eq. 18) when the epigenetic variants have fitness intermediate between the genetic wildtype and mutant, i.e., 0 *< α <* 1; *x* = *b, B*. (a) Only beneficial epimutations occur 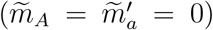 with 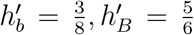. (b) The epigenetic rates depend only on the underlying ge-netic alleles 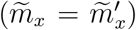 with (i) 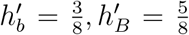 and (ii) 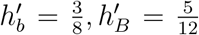. (c) The epigenetic rates depend only on the environment (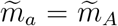 and 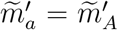) with (i) 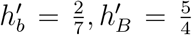 (ii) 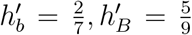 and (iii) 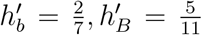. Other parameters: *µ* = *ν* = 10^−6^, *s* = 0.01, *s*′ = 0.02, *r* = 0.01, *h*_*B*_ = 0.1, *h*_*b*_ = 0.2.

Of course, epimutation may not always occur in a beneficial direction. At one extreme, epimutation rates could be completely determined by the features of an allele, 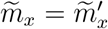. For example, CpG sites are known to be more prone to methylation than other sites (Jaenisch and Bird, 2003). When the epimutation rates are allele-specific and environment-independent and the fitness of the epigenetic variants is intermediate, increasing effective epimuation rates have contrasting effects on rescue. Increasing the effective epimutation rate of the mutant supplies more genetic variation (Fig. S4b), which helps rescue, but also lowers its establishment probability (Fig. S3b), which hurts. The net effect is a reduction in rescue probability as we increase epimutation of the mutant (different colors in Fig. 7b). On the other hand, increasing the effective epimutation rate of the wildtype is always beneficial for rescue (Fig. 7b) since it leads to more *de-novo* mutants (Fig. S3b), and also increases the number of mutants in the standing variation (Fig. S4b).

At the other extreme, epimutation rates could be determined solely by the environ-ment, 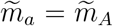 and 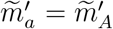. For example, phosphorous starvation in *Arabidopsis* leads to genome-wide hypermethylation irrespective of alleles (Secco *et al*., 2015). Increasing the effective epimutation rate pre-change increases the probability of rescue (Figure 7c) because it increases the number of genetic mutants in the standing variation (Fig. S4c). However, increasing the effective epimutation rate post-change has a non-monotonic effect because it increases number of mutants that arise *de novo* but decreases their individual probabilities of establishing (Fig. S3c). Initially, the cost outweighs the benefit and increasing 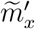 reduces the probability of rescue. In fact, if the genetic threshold is less than the epigenetic threshold, 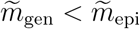, then the rescue probability decreases to zero on reaching the genetic threshold and stays at zero until the epigenetic threshold, after which the rescue probability rapidly increases to 1 (Fig. 7cii). In contrast, if the epigenetic threshold is lower than the genetic threshold, the rescue probability never reduces to zero (Fig. 7ci). In this case, the rescue probability starts to increase even before reaching the epigenetic threshold as the increase in the number of *de novo* mutations can compensate for the decrease in their establishment probability. Finally, if the epigenetic threshold, 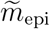, is greater than 1, then rescue is impossible via epigenetics alone (Fig. 7ciii).

### Diploid selection

With diploid selection, rescue by epigenetics is analogous to the case in haploids (Eq. 11), with

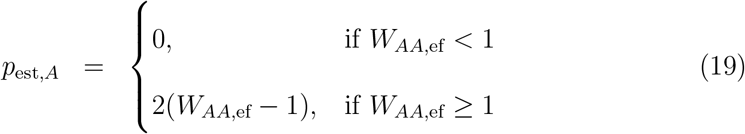

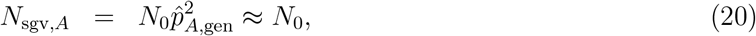

where the equation for *N*_sgv,*A*_ assumes the mutant is rare. Assuming the genetic heterozygote is supercritical, *W*_*Aa*,ef_ *>* 1, the probability a rare genetic mutant rescues the population is then as in the haploid case (Eq. 14) with

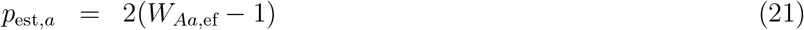

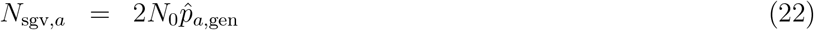

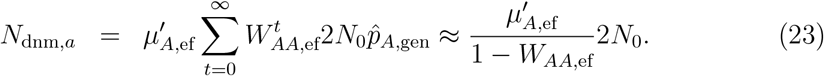

The total probability of rescue is given by Eq. 18.

In our model, a haploid genotype has only two variants (e.g., genetic allele *A* is associated with unmodified *A* and the modified *B*) so their effective fitness is a monotonic function of the QEE. Consequently, altering modification and reset rates has a monotonic effect on each of the survival factors described above (Figs. S3-S4). Since altering the rates changes at most two factors, this can create at most a single maxima or minima in the rescue probability (Fig. 7). In contrast, in diploids, each genetic homozygote is associated with three variants and the genetic heterozygote is associated with four (Fig. 8e,f). This allows for non-monotonic effects on individual survival factors: the number of *de-novo* mutants (Fig. 8b), the number of mutants in the standing genetic variation (Fig. 8c), and the establishment probability of a single mutant (Fig. 8d). Since altering the QEE can influence more than one factor, this can lead to more complex patterns in the probability of rescue (Fig. 8a).

**Figure 8.**
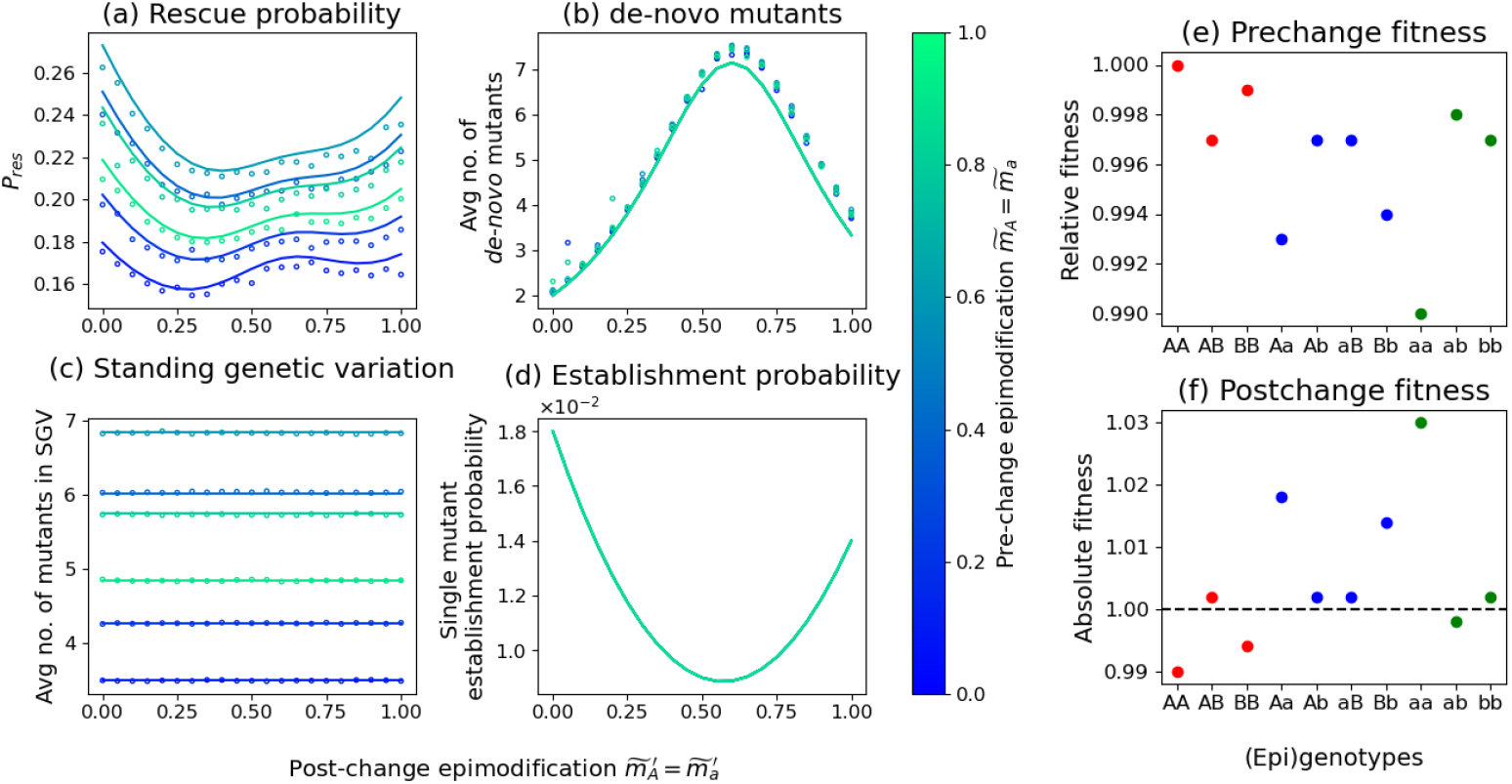
The probability of evolutionary rescue with epigenetics in diploids. (a) The probability of evolutionary rescue in diploids along with the key components, namely the number of mutants from (b) *de-novo* mutations and (c) standing genetic variation, and (d) the establishment probability, plotted as functions of the post-change QEE (identical for both alleles) for different values of the pre-change QEE (different colors). Lines are analytical approximations (Eqs. 18, 21–23) and dots are simulations. (e) Pre-change relative fitness and (f) post-change absolute fitness for each (epi)genotype, grouped by underlying genotype (different colors). Parameter values: *s* = 0.01, *s*′ = 0.02, *r* = 0.01, *µ* = *ν* = 10^−6^.

### Phenotypic plasticity can help or hinder rescue

The effect of phenotypic plasticity on adaptation is a longstanding question in evolutionary biology. Recently, Lambert *et al*. (2025) made a key contribution to this debate from the perspective of evolutionary rescue using a one-locus haploid population genetic model where the wildtype allele switched to an adaptive phenotype in the new environment with probability *p*. Ignoring standing genetic variation, they showed that low levels of plasticity, *p*, can delay extinction of the wildtype (weak Baldwin effect), intermediate levels of plasticity can aid population survival even without genetic adaptation (strong Baldwin effect), and high levels of plasticity can reduce the establishment of genetic mutants by increasing competition with the wildtype (Mayr’s effect). Ignoring competition, our model reduces to theirs when we disregard standing variation, only the wildtype can be modified 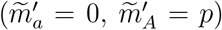 and the epigenetic variant is beneficial (*h*_*B*_ *>* 0). By comparing the probability of rescue with and without plasticity (epigenetics), we see that our model recovers the weak and strong Baldwin effects (light and dark red regions of Fig. 9a, respectively; compare to Fig. 6 of Lambert *et al*. 2025). We can also relax the assumption of no standing variation and no mutant plasticity, which highlights two novel effects of plasticity: (i) plasticity in the pre-change environment can alter standing variation and (ii) plasticity in the post-change environment can alter the mutant’s establishment probability. For instance, in Fig. 9b, pre-change plasticity increases the frequency of genetic mutants in the standing variation while post-change plasticity in the mutant reduces its establishment probability. Whether the net effect is positive or negative depends on the strength of plasticity and the genetic mutation rates. Therefore, whether epigenetics and, more broadly, plasticity aids evolutionary rescue depends on the interplay of multiple effects, including but not limited to the effects proposed by Baldwin (Baldwin, 1896; Simpson, 1953) and Mayr (Mayr, 1970).

**Figure 9.**
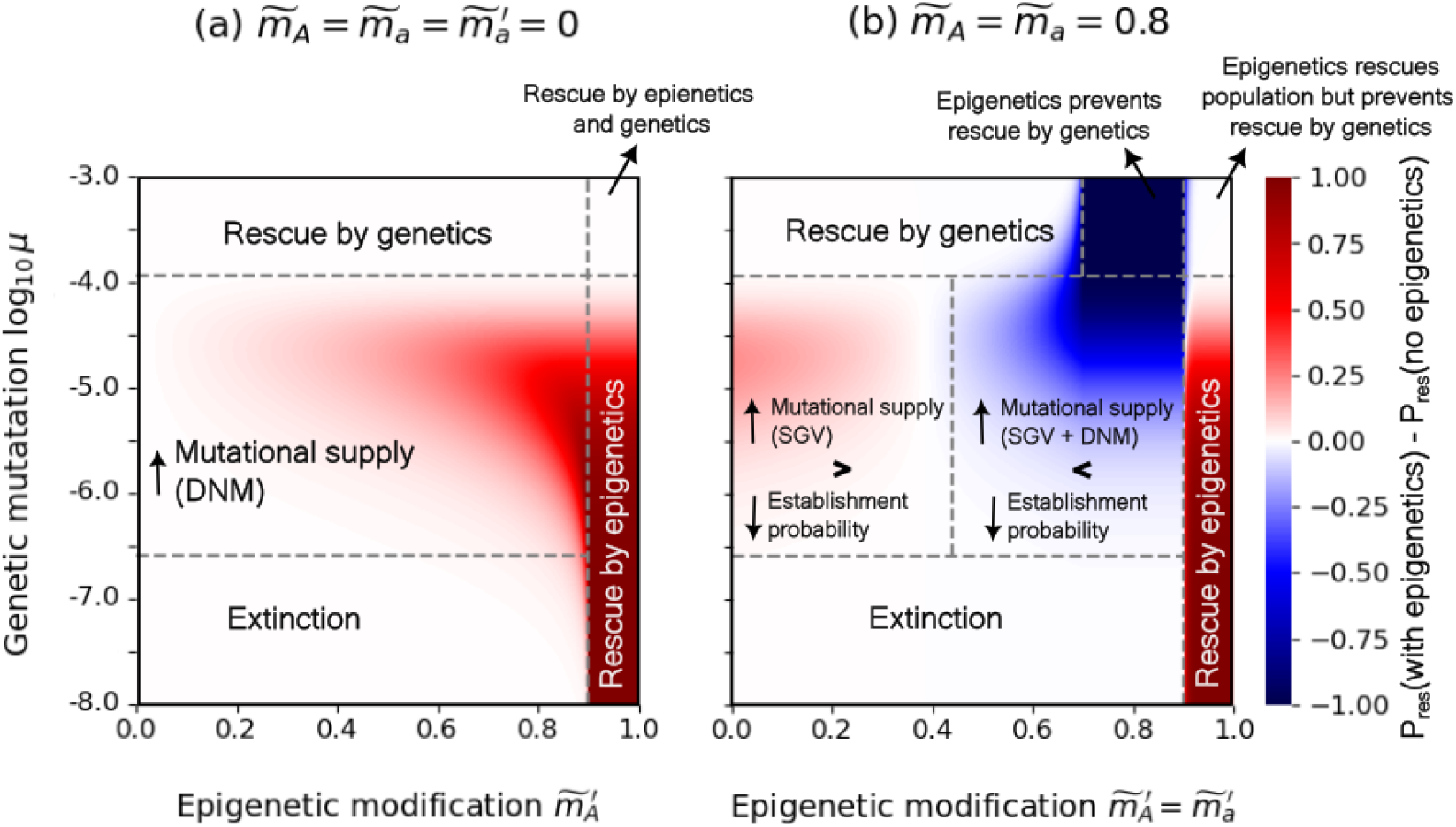
The effect of epigenetics on evolutionary rescue. Epigenetics increases the probability of rescue in the red regions and decreases it in the blue regions. This is plotted across the post-change QEE of *A*, 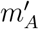, and genetic mutation rates, *µ* = *ν*. The regions labeled “rescue by genetics” refers to the parameter space where rescue by the establishment of the genetic mutant is high, irrespective of epigenetic dynamics, and “rescue by epigenetics” refers to regions where rescue is more likely to occur by the establishment of the genetic wildtype mediated by the epigenetic variant, increasing its effective fitness. (a) Only post-change epigenetic modification of *A*, similar to Lambert *et al*. (2025). (b) Allowing for epigenetic variants of both alleles, with epigenetic rates determined by the environment. The rescue probability in both figures is calculated from Eq. 18. The horizontal lines in both panels are the mutation rate at which the rescue probability without epigenetics is 1% and 99%, respectively (Eq. S8). The rightmost vertical line in both panels delineating the epigenetic rescue region is 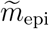. The other two vertical lines in panel (b) are the thresholds when epigenetics become detrimental relative to the case with no epigenetics (Eq. S9) and 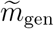. See Section S4 for more details. Other parameters: 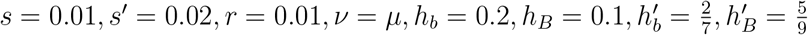.

## Discussion

Epigenetics can affect evolution, with wide-ranging consequences for antibiotic and pathogen resistance (Ghosh *et al*., 2020), cell state heterogeneity in cancer (Davalos and Esteller, 2023), aging in vertebrates (Pal and Tyler, 2016), and the maintenance of phenotypic variation (Duncan *et al*., 2014), amongst others Ashe *et al*. (2021). A major obstacle to better understanding the role of epigenetics is the inherent complexity of incorporating more variables and parameters into evolutionary models, prevent-ing a general analysis. Here we show how to reduce the complexity of population epigenetic models to their underlying population genetic model by introducing the quasi epigenetic equilibrium (QEE), which is valid whenever the epigenetic dynamics happen fast relative to genetic mutation, selection and drift (Fig. 2a). This often simplifies the model to one that is already well-studied, producing immediate insight. For example, here we show that models of one-locus haploid and diploid selection with epimutation, which have four epialleles to track, reduce to their respective two-allele classical forms under the QEE. Standard mutation-selection balance results then inform how minor changes in epigenetic dynamics – which affect the parameters of the underlying genetic model – can lead to drastic changes in the evolutionary dynamics (Fig. 2b and Fig. 3a). Further, we can understand the role of epimutation on population survival after an abrupt change in the environment by combining the QEE with existing evolutionary rescue results, which shows that epigenetics generally has a nonmonotonic effect on survival, aiding or impeding it (Figs. 7, 8, 9). We also show how the QEE extends to more complex scenarios; a one-locus model with paramutation, where epimutation only happens in heterozygotes, reduces to a one-locus two-allele model with frequency-dependent selection (Fig. 4).

### Epigenetics can affect evolution without being inherited

The form of the QEE itself provides some key insights into the important properties of epigenetics from an evolutionary perspective. In particular, the leading-order QEE derived above, 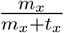 (Eqs. 4 and 9), depends only on the ratio of epigenetic transition probabilities and not on their absolute values. This ratio can be reasonably large even with guaranteed reset, *t*_*x*_ = 1, as long as modifications occur frequently (high *m*_*x*_). This implies that epigenetic modifications need not necessarily be inherited to have a significant evolutionary effect. Frequent modifications can arise for a number of reasons, including a consistent recurrent response to the environment (Bogan and Yi, 2024; Dowen *et al*., 2012), proximity to epigenetically silenced transposable elements (Choi and Lee, 2020), or an allele that regularly recruits epigenetic modifications (Issa, 2004). Consistent but non-heritable variation, also known as phenotypic variability or bet-heging, is known to affect evolution (Childs *et al*., 2010) and the role of epigenetics in mediating it has also been demonstrated, for instance, in yeast (Bódi *et al*., 2017), as well as other systems (Stajic and Jansen, 2021).

### Epigenetic-mediated shifts in selection, dominance and mutation

The magnitude of the QEE is consequential because it can alter any parameter of the reduced model. Perhaps most obviously, the QEE determines the effective fitness of the genetic alleles, thereby influencing the fitness landscape. For instance, in the haploid model we see that increasing the value of the QEE reduces the effective fitness difference between the two genetic alleles (i.e., smoothens the fitness landscape) when the epigenetic variants have fitness intermediate to that of the unmodified genetic alleles (0 *< h*_*x*_ *<* 1) (as in Mall *et al*., 2025). This is, of course, not necessarily the case; if the epigenetic variants do not have intermediate fitness, then increasing the QEE can exacerbate fitness differences (i.e., make the fitness landscape more rugged). Further, in diploids, even when the epigenetic variants have intermediate fitness (0 *< h*_*xy*_ *<* 1), the QEE can roughen the fitness landscape through dominance. For example, in Fig. 3a, increasing the QEE switches the reduced system from directional selection, to overdominance to underdominance, with huge effects on the evolutionary equilibrium. The effect of epigenetics on dominance may be particularly interesting when epigenetic rates change with the environment. The resulting epigenetic-mediated shifts in dominance are then predicted to have a number of downstream consequences, e.g., on the benefit/cost of sex in evolutionary rescue (Uecker, 2017), the softness of selective sweeps (Muralidhar and Veller, 2022), and the stabilization of polymorphisms in seasonal environments (Brud, 2025). Also, since dominance in diploid models is analogous to epistasis in haploid models (Otto, 2003; Uecker, 2017), our results imply that there can be epigenetic-mediated shifts in epistasis, with consequences for the role of recombination in evolutionary rescue (Uecker and Hermisson, 2016) and trade-offs between alternative evolutionary pathways (Hall *et al*., 2019). The QEE can also influence the effective mutation rate, potentially resulting in stress-induced mutagensis, which can provide mutational supply in novel environments without the detrimental effects of high mutation rates in stable environments (Matic, 2019).

### The multi-faceted effects of epigenetics on evolutionary rescue

The fact that the QEE is reached within a few generations means it is not only useful for capturing long-term effects, like mutation-selection balance, but also phenomena that occur on rapid timescales, like evolutionary rescue. This allows us to address the related question of whether plasticity is beneficial for adaptation and persistence.

Here, facilitated by the QEE, we find that epigenetics can alter the probability of evolutionary rescue in four distinct ways: by affecting (i) the number of mutants in the standing variation, (ii) the number of mutants that arise during the decline of the wildtype, (iii) the establishment probability of a mutant and (iv) whether the wildtype can sustain the population on its own. Whether the net effect of epigenetic-driven changes in these factors is positive or negative depends on (i) the fitness of the epigenetic variants, (ii) the correlation in epigenetic transition probabilities between alleles and environment and (iii) the effect of epigenetics on other evolutionary parameters, such as genetic mutation rates. This shows that epigenetic rates can have complex effects on the probability of evolutionary rescue (Fig. 8), facilitating or hindering survival (Fig. 9).

The effect of epigenetics on population persistence is increasingly documented and varied (Mounger *et al*., 2021; Stajic and Jansen, 2021). For instance, yeast populations can develop resistance to an acid via epigenetic silencing (Stajic *et al*., 2019). Higher epigenetic silencing rates slow the decay of the sensitive strains, allowing genetically-resistant mutants to arise, though the epigenetic mechanisms also reduce the fitness of these genetic mutants. Similarly, in green algae, epigenetic variation increases initial growth rates and the number of genetic mutations that accumulate, either by increasing the effective mutation rate or decreasing the decline rate of the susceptible population, in a variety of stresses (Kronholm *et al*., 2017). In contrast, the epigenetic response of*Daphnia*to an environmental stressor (the cyanobacteria*Microcystis*), which increasesfitness variance, appears to be maladaptive (Shahmohamadloo *et al*., 2025). While evidence of survival purely by epigenetics is limited, Japanese knotweeds provide a compelling example, as genetically identical strains have been able to successfully invade different regions of central Europe, with epigenetic variation being correlated to local environments (Zhang *et al*., 2016). Using the QEE, we were able to analyze a very general model of epigenetics in a changing environment, capturing a diversity of observed epigenetic responses.

### Future directions

The analytical tractability provided by the QEE now allows us to investigate the role of epigenetics in more complex evolutionary scenarios. For instance, epigenetic heterogeneity amongst cells within an individual is common (Carter and Zhao, 2021; Webster and Phillips, 2025). If the epigenetic state an offspring inherits from its parents completely determines the epigenetic variation across its cells, and thus its fitness, then the models we have analyzed above apply directly. However, if there is stochasticity in epigenetic variation across cells for a given inherited state, then we need a more complex model (Biwer *et al*., 2020; Flanagan *et al*., 2006). In either case, the QEE could be useful in determining the within-organism distribution of epigenetic states, as long as the strength of selection between cells is weaker than within-individual epigenetic modification rates. We could then potentially analyze the system by combining the QEE with classical multilevel selection theory (Okasha, 2006).

Another application is understanding the emergence of plasticity influctuating environments via the evolution of epigenetic modification rates. A recent study (Romero-Mujalli *et al*., 2024) investigated this, assuming that epigenetic modifications are purely beneficial and affect a single trait. Our method provides an analytical approach to generalize this study to modifications of arbitraryfitness effects and multiple traits, allowing a more complete picture of whenfluctuating environments promote the evolution of epigenetic modifications.

Our primary focus here was the effect of epigenetics on genetic evolution and population persistence. Wefind that epigenetics influences both primarily by determining the effectivefitness of genetic alleles. However, we can also look at how epigenetics affects phenotypic evolution, which is determined by selection, heritability, variation, and transmission bias (Okasha, 2006). In a preliminary investigation (Section S5), we find that epigenetics can alter all of the components of phenotypic evolution. More importantly, the QEE and effective parameters of genetic alleles (fitness, mutation, and phenotype) are insufficient. In particular, the phenotypic distinction between different forms of the same genetic allele plays a major role, e.g., we need to consider phenotypic variation both between and within genetic alleles. Future work in this vein could explore the downstream effects of epigenetics on phenotypic evolution, such as evolutionary rescue via a quantitative trait.

## Data availability statement

All simulation code and data is available at https://github.com/pderaje/Epigenetic_Rescue. Upon publication of the article, it will be made available on Zenodo. The figures were made in jupyter-notebook named `FinalFigures.ipynb` and the calculations are in a mathematica file titled `PopEpi Mathematica.nb`, both can be found in the github folder.

## Acknowledgements

We thank Aneil Agrawal, Ina Anreiter, Marla Sokolowski, Stephen Wright, and the Osmond lab for helpful conversations. This work was supported by the Natural Sciences and Engineering Research Council of Canada (RGPIN–2021-03207 and DGECR-2021-00114 to MO). Computations were performed on the Niagara supercomputer at the SciNet HPC Consortium. SciNet is funded by: the Canada Foundation for Innovation; the Government of Ontario; Ontario Research Fund – Research Excellence; and the University of Toronto.

## S1 Standard epimutation

### First order QEE

The zeroth-order QEE with standard epimutation is given in the main text (Eqs. 4a–4b). We can also calculate the QEE tofirst order,

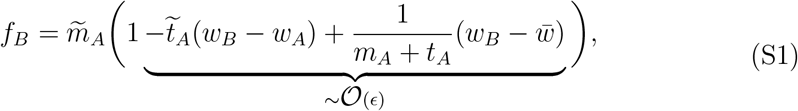

with an analogous equation for *f*_*b*_, where 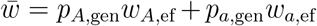 and the effective fitnesses are calculated using the zeroth-order approximation of *f*_*B*_ and *f*_*b*_. Note that the first-order term depends on allele frequency, *p*_*A*,gen_, via population mean fitness, 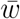, but this only has an appreciable effect if *m*_*A*_ + *t*_*A*_ is small, in particular of order 𝒪 (*ϵ*). The QEE is therefore still roughly independent of allele frequency when *m*_*x*_ + *t*_*x*_ *~* 𝒪 (1), which is a weaker condition than assuming both *m*_*x*_ and *t*_*x*_ are 𝒪(1).

## S2 Paramutation

### Paramutable mutant, special case

When the mutant *a* is paramutable, its epiallele *b* has the same fitness as the wildtpye *A*, and the unmodified mutant allele *a* is recessive, we need to use the second-order approximation (in *ϵ*) of the equilibrium allele frequency,

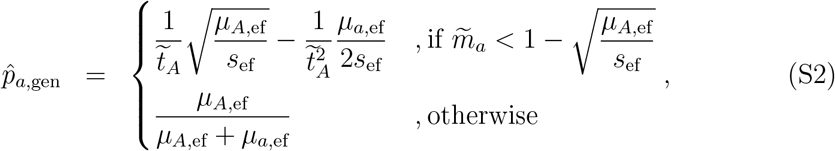

where *s*_ef_ = *w*_*AA*,ef_ −*w*_*AA*,ef_.

### Paramutable wildtype, first-order QEE and allele frequencies

When the wildtype *A* is paramutable the first-order QEE is

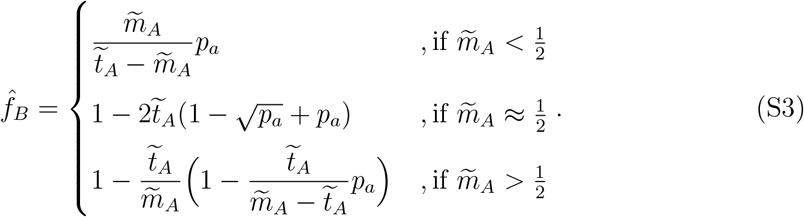

Substituting this QEE into the effective fitnesses and mutation rates, we get a frequencydependent two-allele model. Solving this we get the equilibrium frequencies for the genetic mutant at mutation-selection balance. This is not possible in general but it is for special cases. In particular, assuming *B* has the same fitness as *a* and writing *h* = *h*_*Aa*_ = *h*_*aB*_, when the mutant is recessive (*h* = 0) its equilibrium frequency is given by

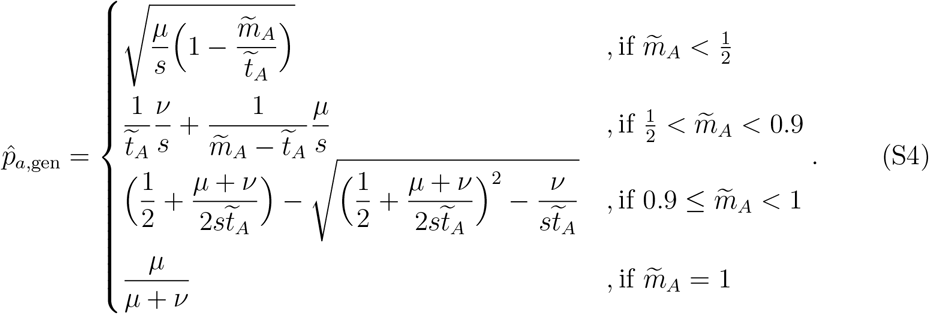

When the mutant is not recessive its equilibrium frequency is

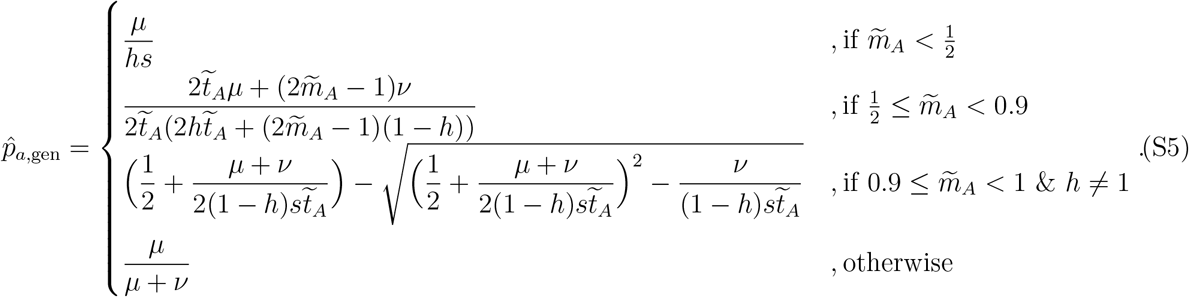

Note that when 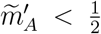, the result for *h ?*= 0 is consistent with using Eq. 9 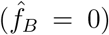 in Eq. 7 but this is not the case for a recessive mutant, *h* = 0, where the first-order QEE term alters the equilibrium frequency (with 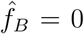 and *h* = 0, Eq. 7 gives 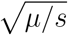). To understand this, note that when the mutant is not completely recessive (*h ?*= 0), the equilibrium frequency of the mutant *a* is not altered by weak paramutation (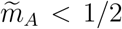; Fig. 4e,f). Once the threshold 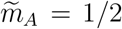 is crossed, we see a rapid increase in the frequency of both the mutant *a* and the epigenetic variant *B*. This is due to the frequency-dependent effect of paramutation; the *B* epiallele, despite being deleterious, can spread due to its transmission advantage over wildtypes in heterozygotes. This is similar to gene-drive, except the transmission advantage is reduced by reset. This transmission bias also reduces the effective fitness of the wildtype *A*, which increases the relative fitness of the mutant *a*, allowing it to reach higher frequencies. When the mutant is dominant, the marginal fitness of genetic allele *A* reduces significantly, which can accelerate the feedback between paramutation and epiallele frequency. We then need to take into account the first-order terms in the QEE to get good approximations at high values of 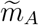(Fig. 4f). However, when the mutant is completely recessive (*h* = 0), mutant frequency declines with 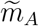 before it begins to increase (Fig. 4d). This is because, even when rare, paramutation in heterozygotes exposes mutants to selection that they would otherwise be shielded from. The detriment from this outweighs the benefits obtained from the transmission advantage and the reduction in the fitness of the wildtype at low paramutation rates. At higher paramutation rates, the benefits outweigh the cost and the mutant increases for similar reasons as in the non-recessive case.

## S3 Haploid rescue

### Epigenetics only, near criticality

When there is no genetic variation, i.e., *p*_*A*,gen_ = 1, substituting the post-change fitnesses *W*_*A*_ = 1 −*r* and *W*_*B*_ = 1 −*r* + *s*′ into the first-order QEE (Eq. S1) gives

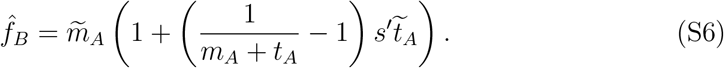

Subbing this into effective fitness, *W*_*A*,ef_, near criticality, 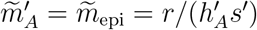, gives

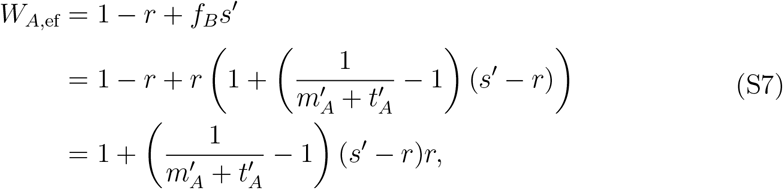

which decreases with 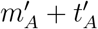.

## S4 Details of Fig. 9

The dashed lines in Fig. 9 were computed as below:

- Horizontal lines are the genetic mutation rates such that the rescue probability in the absence of epigenetics is 1% and 99% respectively. This obtained by

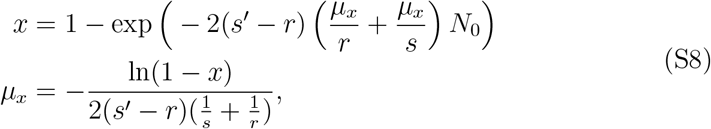

where *x* = 0.01 and *x* = 0.99, respectively.
- The rightmost vertical line is the threshold after which the genetic wildtype is supercritical, given by 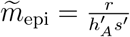.
- The leftmost vertical line in panel b is obtained by solving for the post-change 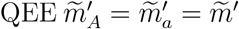 that makes the product of mutational supply from standing variation and establishment probability the same in the two cases (with and without epigenetics),

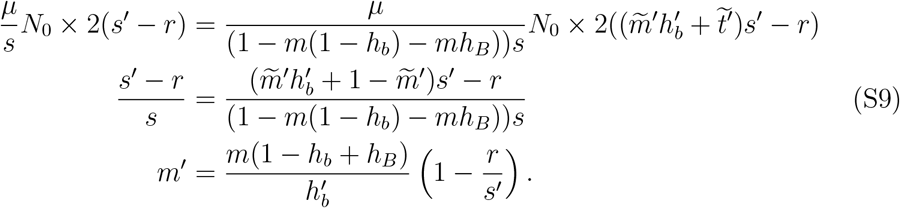
- The middle vertical line at the top of panel b is the post-change QEE that makes the establishment probability of the genetic mutant zero,

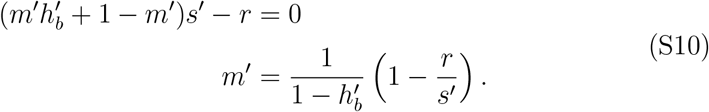

This is 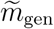.

## S5 Effect of epigenetics on genetic and phenotypic evolution

The change in the mean value of a character over one generation is given by

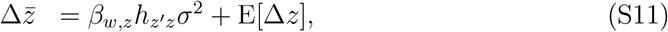

where *β*_*wz*_ is the selection gradient (the slope of the regression of fitness, *w*, on trait value, *z*), 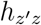 is the heritability (the regression coefficient of the mean offspring trait value, *z*′, on the parent value, *z*), 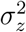 is the variance in parent trait values and E[Δ*z*] = *∫*_*z*_(*z*′−*z*)*dz* is the transmission bias (Eq.1.3 of Okasha, 2006).

Now, consider a haploid one-locus two-allele model without epigenetics. Let the phenotype of an individual with allele *x* be *z*_*x*_ and their offspring’s expected value be 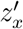. Then we have

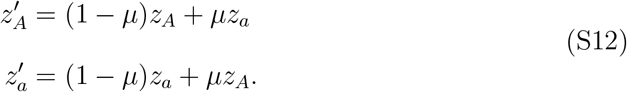

and the components contributing to phenotypic evolution in Eq. S11 are

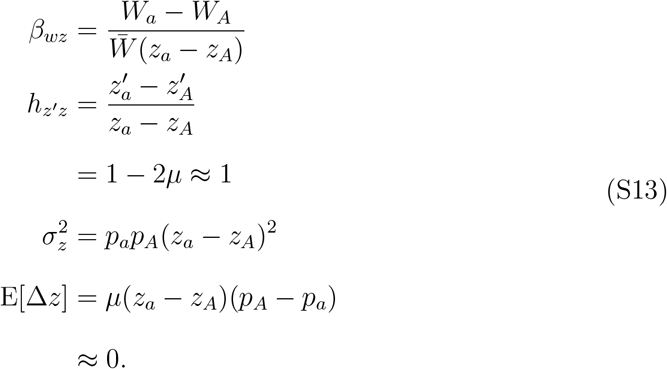

Plugging in *z*_*a*_ = 1 and *z*_*A*_ = 0, to consider the change in the frequency of the mutant allele,

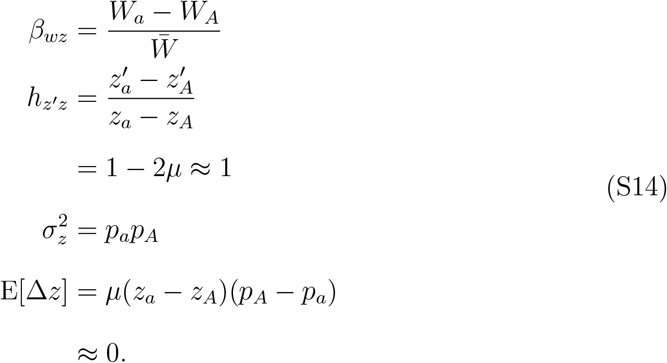

This gives the classic equation for allele frequency dynamics, 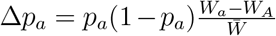.

With standard epimutation we have

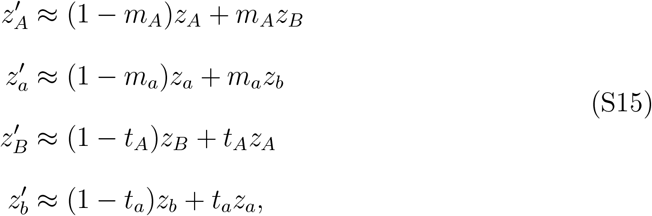

which gives

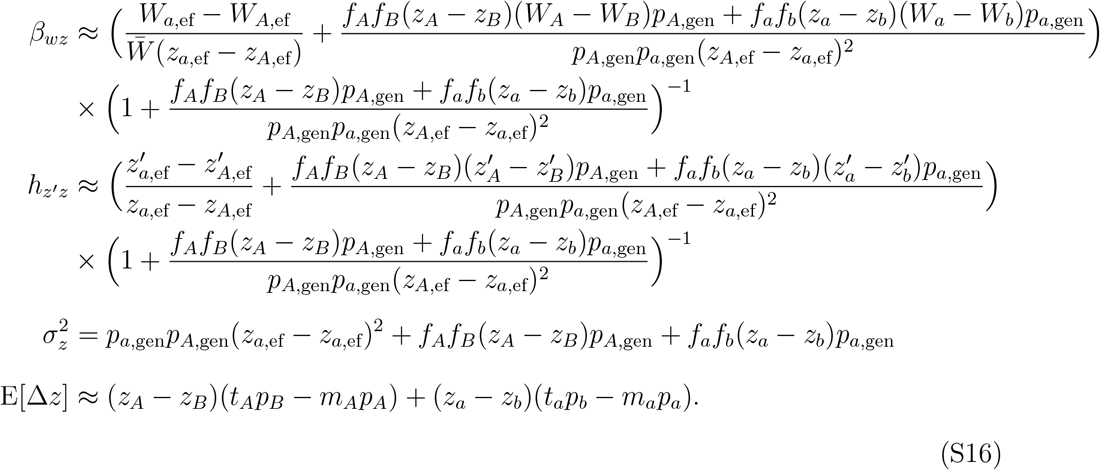

From this we see that epigenetics can affect all aspects of phenotypic evolution: selection, heritability, and variation. Further, there are deviations from what is obtained by simply substituting the effective fitnesses and effective phenotypes into the previous equations without epigenetics. These deviations depend on the phenotypic effects of modifications, *z*_*A*_ −*z*_*B*_ and *z*_*a*_ −*z*_*b*_. However, if we now consider the effect of epigenetics on genetic evolution (by putting *z*_*A*_ = *z*_*B*_ = 0 and *z*_*a*_ = *z*_*b*_ = 1), we get

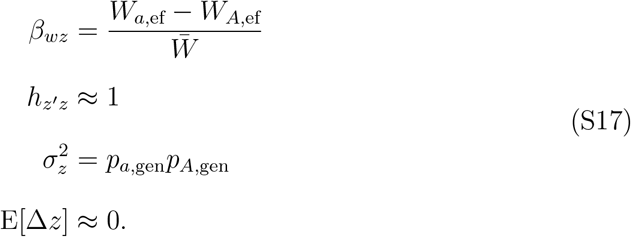

Therefore, epigenetics affects genetic evolution only by altering the fitness of genotypes, which can be captured by the QEE-weighted effective fitnesses, 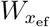.

## S6 Supplementary figures

**Figure S1:**
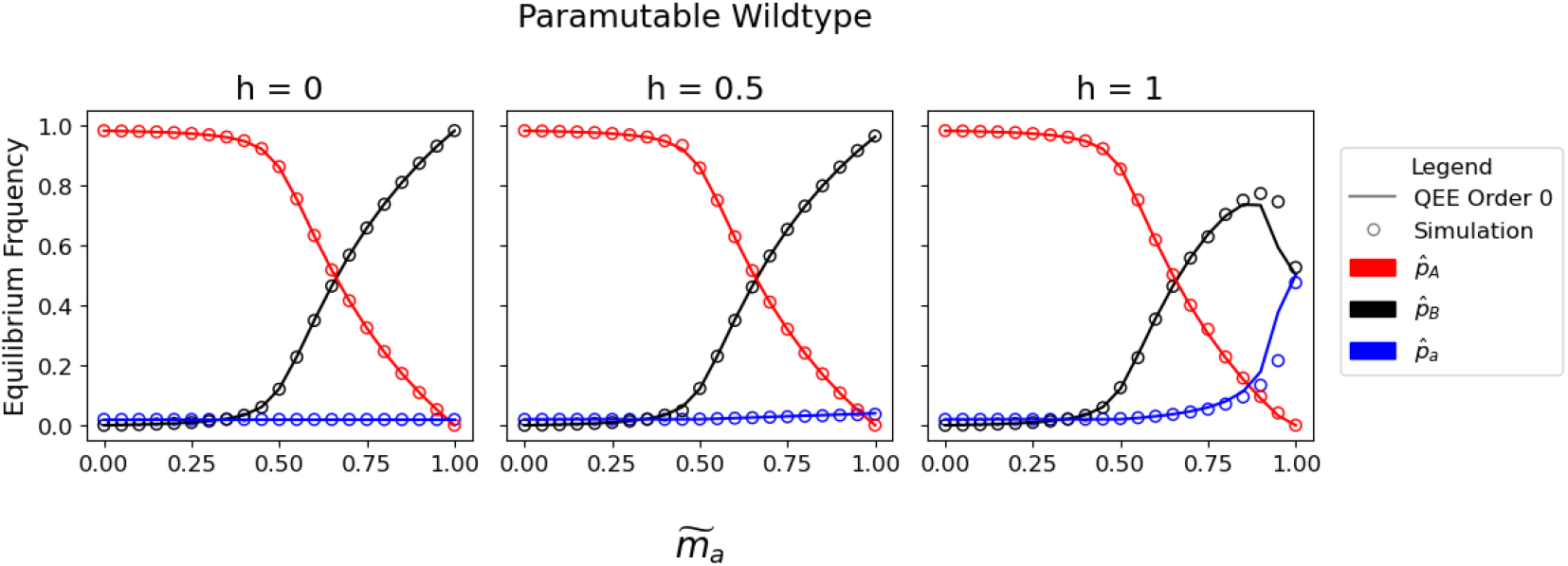
The effect of paramutation on equilibrium allele frequencies. The wildtype allele *A* is paramutable. In contrast to Fig. 4d-f all (epi)heterozygotes have fitness exactly midway between the two (epi)homozygotes: 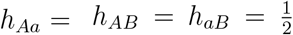. The empty circles are from deterministic simulations while the lines are analytical approximations (Eq. 7 with 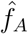 from Eq. 9). Other parameters: *µ* = *ν* = 10^−4^, *s* = 0.01.

**Figure S2:**
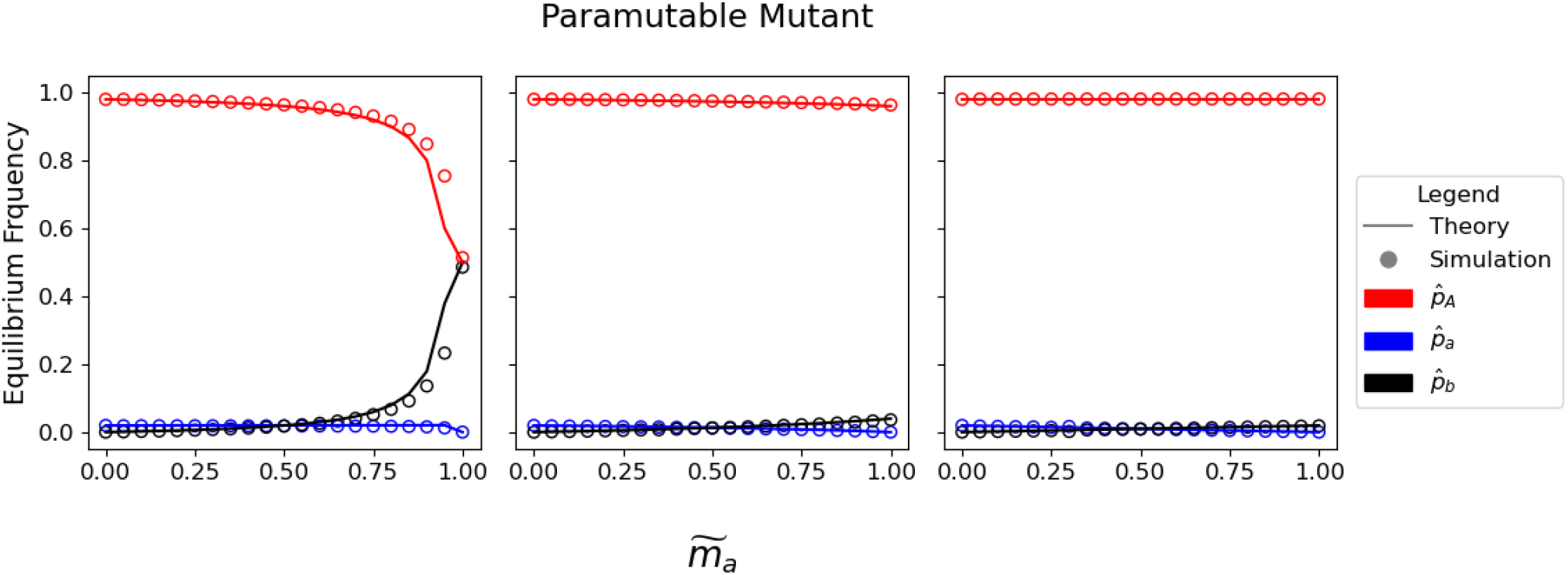
The effect of paramutation on equilibrium allele frequencies. The mutant allele *a* is paramutable. In contrast to Fig. 4a-c, all (epi)heterozygotes have fitness exactly midway between the two (epi)homozygotes: *h*_*bb*_ = *h* and 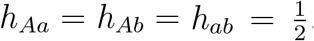. The empty circles are from deterministic simulations while the lines are analytical approximations (Eq. 7). Other parameters: *µ* = *ν* = 10^−4^, *s* = 0.01.

**Figure S3:**
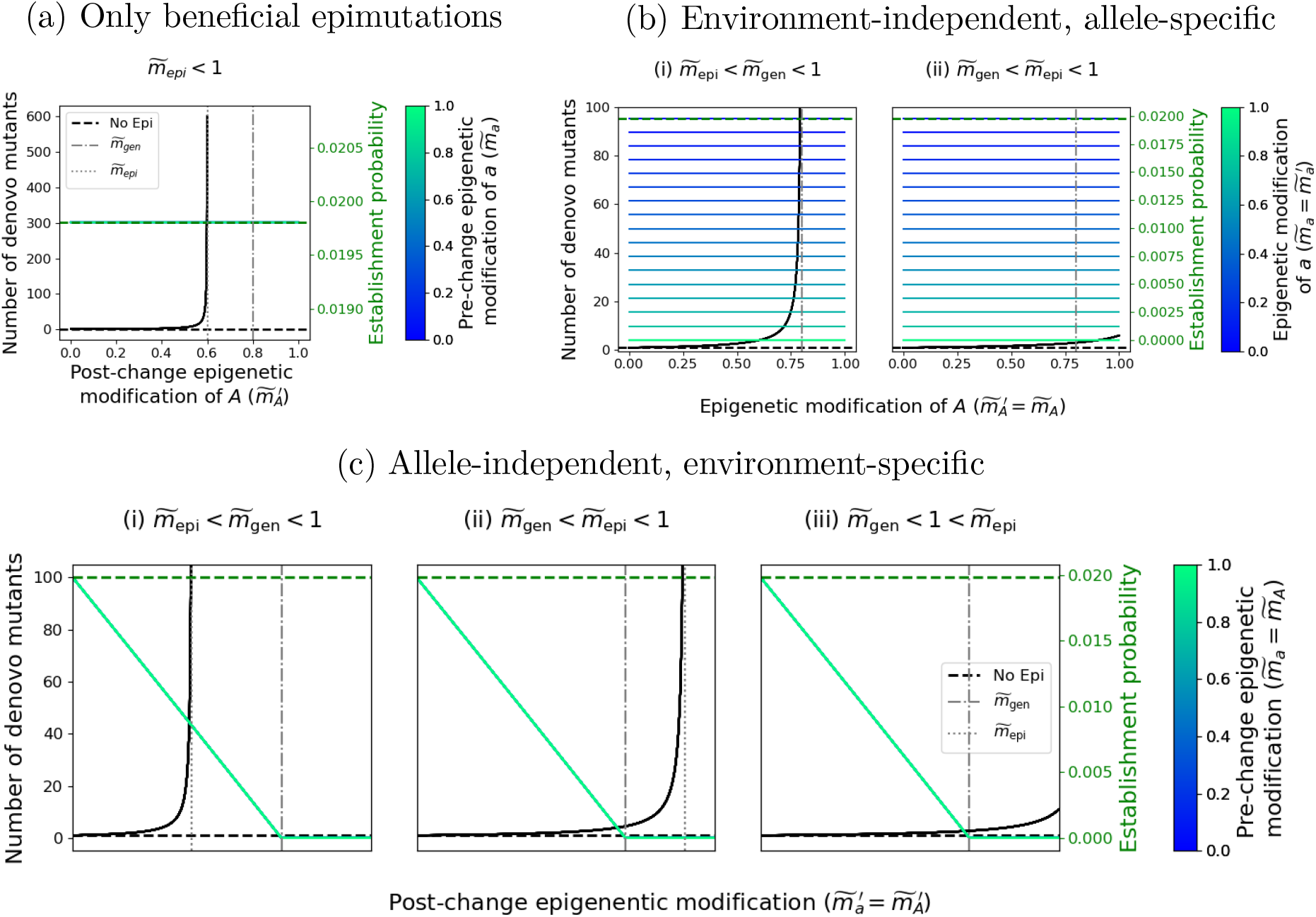
Effect of epigenetics on the number of *de-novo* mutations and their establishment probability. Number of *de novo* mutants (black lines) and the establishment probability of a single mutant (blue-green lines) for the haploid model when epigenetic variants have fitness intermediate between the genetic wildtype and mutant, i.e., 0 < *h*_*x*_ < 1; *x* = *b, B*. The parameters correspond to Fig. 7. The dashed lines are the respective values when there are no epigenetics. (a) Only beneficial epimutations occur 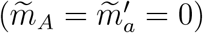 with 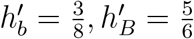. (b) The epigenetic dynamics are insensitive to environment 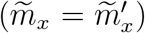 with (i) 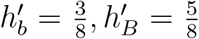 and (ii) 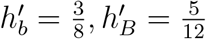.(c) The epigenetic dynamics are only dependent on the environment (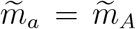 and 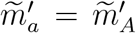) with (i) 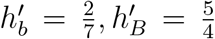, (ii) 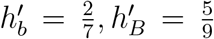 and (iii) 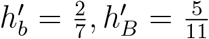. Other parameter values for all panels: *µ* = *ν* = 10^−6^, *s* = 0.01, *s*′ = 0.02, *r* = 0.01, *h*_*B*_ = 0.1, *h*_*b*_ = 0.2.

**Figure S4:**
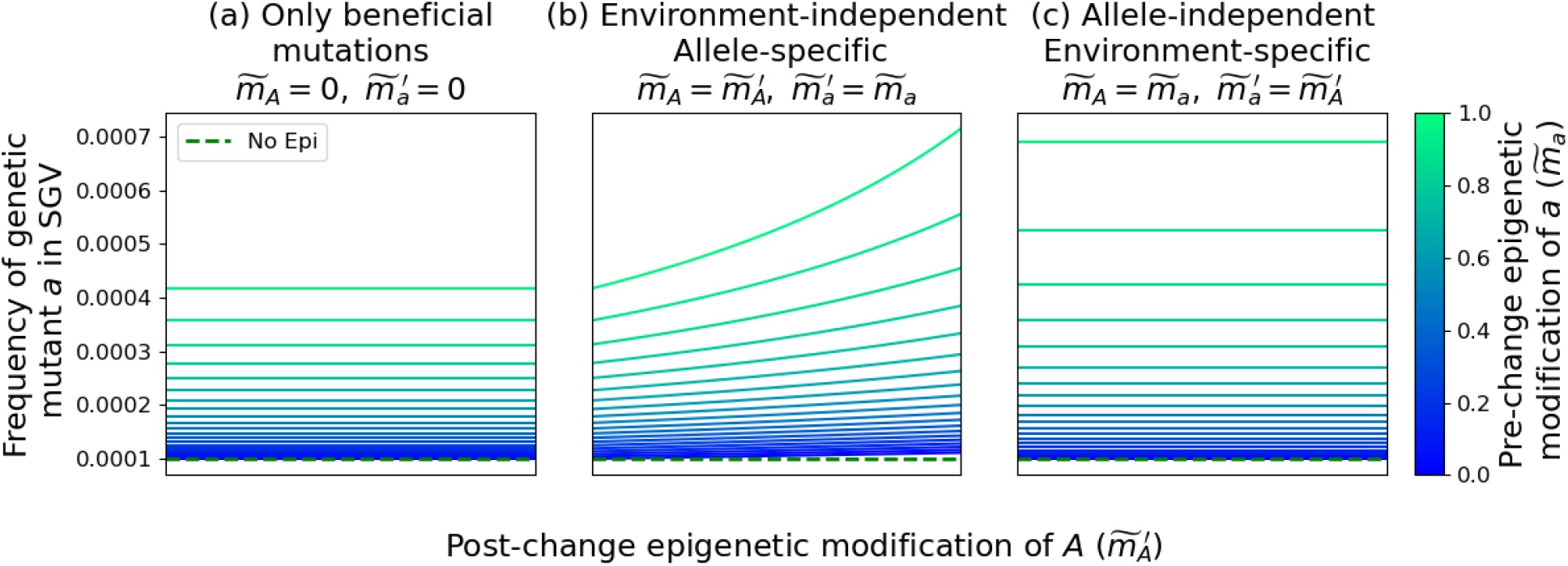
Effect of epigenetics on the standing genetic variation. Frequency of genetic mutants in the standing genetic variation for the haploid model when the epigenetic variants have fitness intermediate between the genetic wildtype and mutant, i.e., 0 *< h*_*x*_ *<* 1; *x* = *b, B*. See Fig. S3 for more information.

